# Post-death Vesicles of Senescent Bone Marrow Mesenchymal Stromal Polyploids Promote Macrophage Aging and Breast Cancer

**DOI:** 10.1101/2024.03.06.583755

**Authors:** Bowen Xie, Ming Fan, Charles X. Wang, Yanhong Zhang, Shanxiu Xu, Rachel Mizenko, Tzu-yin Lin, Yixin Duan, Yanyan Zhang, Jie Huang, Jonathan I. Berg, Douglas Wu, Anna Li, Dake Hao, Kewa Gao, Yaohui Sun, Clifford G. Tepper, Randy Carney, Yuanpei Li, Aijun Wang, Qizhi Gong, Magen Daly, Li-En Jao, Arta M. Monjazeb, Fernando A. Fierro, Jian Jian Li

## Abstract

Potential systemic factors contributing to aging-associated breast cancer (BC) remain elusive. Here, we reveal that the polyploid giant cells (PGCs) that contain more than two sets of genomes prevailing in aging and cancerous tissues constitute 5-10% of healthy female bone marrow mesenchymal stromal cells (fBMSCs). The PGCs can repair DNA damage and stimulate neighboring cells for clonal expansion. However, dying PGCs in advanced-senescent fBMSCs can form “spikings” which are then separated into membraned mtDNA-containing vesicles (Senescent PGC-Spiking Bodies; SPSBs). SPSB-phagocytosed macrophages accelerate aging with diminished clearance on BC cells and protumor M2 polarization. SPSB-carried mitochondrial OXPHOS components are enriched in BC of elder patients and associated with poor prognosis. SPSB-incorporated breast epithelial cells develop aggressive characteristics and PGCs resembling the polyploid giant cancer cells (PGCCs) in clonogenic BC cells and cancer tissues. These findings highlight an aging BMSC-induced BC risk mediated by SPSB-induced macrophage dysfunction and epithelial cell precancerous transition.

**SIGNIFICANCE:** Mechanisms underlying aging-associated cancer risk remain unelucidated. This work demonstrates that polyploid giant cells (PGCs) in bone marrow mesenchymal stromal cells (BMSCs) from healthy female bone marrow donors can boost neighboring cell proliferation for clonal expansion. However, the dying-senescent PGCs in the advanced-senescent fBMSCs can form “spikings” which are separated into mitochondrial DNA (mtDNA)-containing spiking bodies (senescent PGC-spiking bodies; SPSBs). The SPSBs promote macrophage aging and breast epithelial cell protumorigenic transition and form polyploid giant cancer cells. These results demonstrate a new form of ghost message from dying-senescent BMSCs, that may serve as a systemic factor contributing to aging-associated immunosuppression and breast cancer risk.

**Graphic Abstract:** Xie et al demonstrate that the polyploid giant cells (PGCs) in the juvenile phase expansion of female bone marrow mesenchymal stromal cells (fBMSCs) can boost neighboring cell proliferation for clonal expansion. However, when fBMSCs enter to the advanced-senescent phase, the dying-senescent PGCs form “spikings” which are then separated into membraned vesicles termed Senescent PGC spiking bodies, SPSBs). The SPSBs carrying fragmented mitochondrial elements and OXPHOS proteins can be phagocytosed by macrophage and breast epithelial cells leading to macrophage aging and breast epithelial protumorigenic transition. The SPSBs are demonstrated to be a new form of post-cell death vesicle from aging BMSCs and may serve as a systemic factor contributing to aging-associated immunosuppression and breast cancer risk.

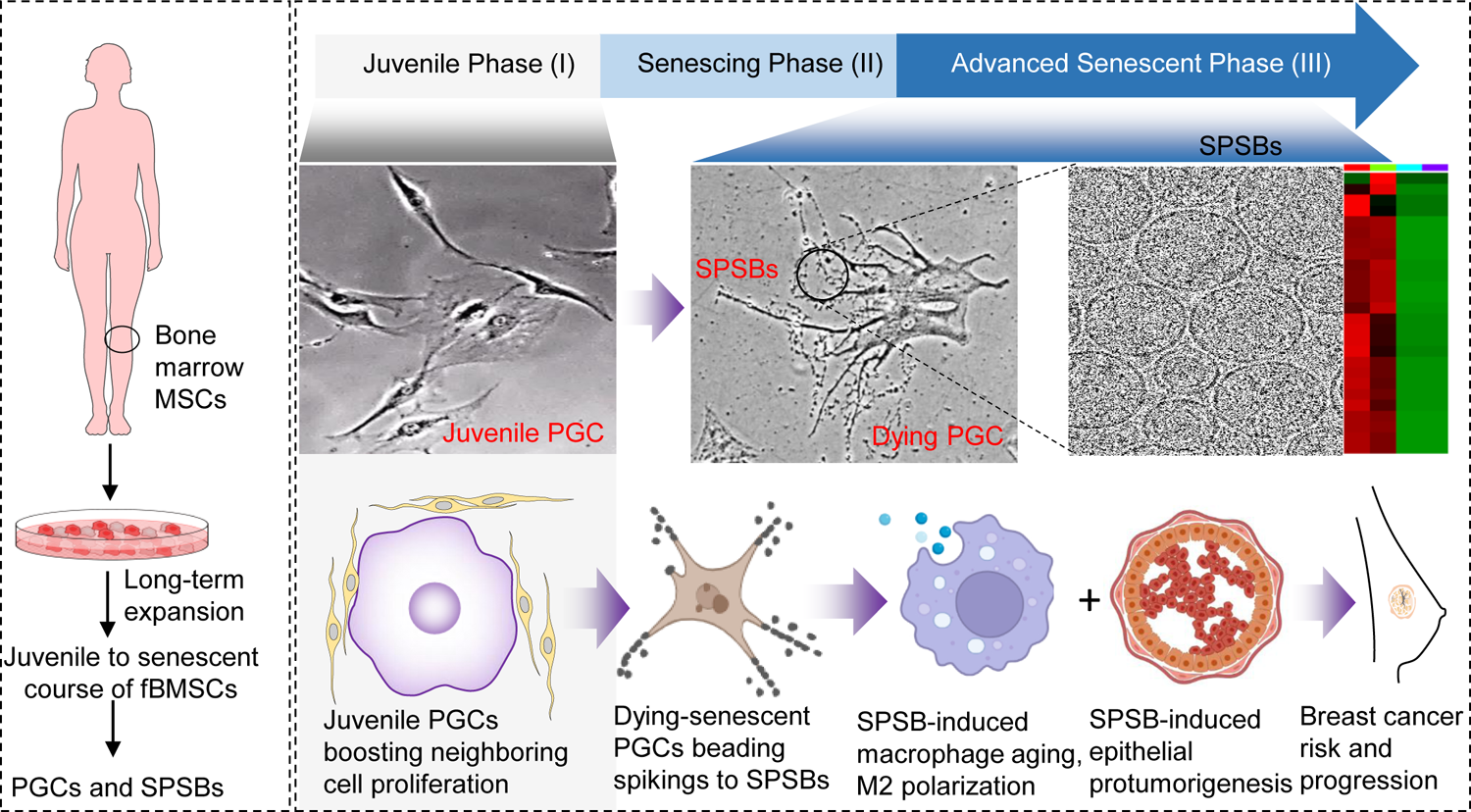

## INTRODUCTION

The mammary gland, an organ subject to dynamic remodeling throughout life, intricately relies on a hemostatic luminal structure sustained by multiple cellular communications. Cell senescence observed by Leonard Hayflick and Paul Moorhead in the 1960s is increasingly evidenced as the aging “hallmark” contributing to aging-associated organ functional decline and cancer risk. Abolishing the senescent cells is extensively investigated for senolytic and senomorphic therapies (1, 2). In aging breasts, senescent stroma-induced macrophage dysfunction and epithelial distortion coordinatively provoke breast cancer (BC) susceptibility and progression (3–7). Beyond the tissue degenerative modifications, aging-induced immune cell suppression including macrophages that are actively involved in both innate and adaptive immune regulations can boost the proinflammatory and protumorigenic breast epithelium (8, 9). In turn, the senescent cells in the aging organ can enhance immune cell exhaustion further accelerating BC susceptibility and tumor aggressive progression(8, 10). The precancerous epithelial structure can impair macrophage functions (6, 11) and promote macrophage pro-tumour polarization and BC metastasis (12). In light of the established milieu of BC risk via coordinative macrophage deficiency and aging-induced breast epithelial distortion, we wondered if potential systemic signals generated from the vast resource of aging stromal tissues could be involved in the aging-associated macrophage suppression and cancer-prone epithelial transition.

In adult humans, bone marrow mesenchymal stromal cells (BMSCs) are an immense supply of stromal cells contributing to an array of essential physiological functions including the multipotent potential of differentiating into diverse types of matured cells and regulation of immune surveillance (13, 14). It is unclear whether a circulating factor could be generated from senescent BMSCs although hematopoietic stem cells can be specifically affected by the BMSC-secreted senescent-associated secretory phenotype (SASP) leading to stem malignant transformation(15, 16). Cargo contents of a fraction of circulating extracellular vesicles (EVs) are indicated to be aging-dependent and BMSC-EVs from young animals are shown to reduce cell senescence and extend the animal life span(17, 18). However, contrasted to the SASP and EVs that are actively synthesized and secreted from senescent yet metabolically active BMSCs, there is a notable scarcity of information regarding the signals emanating from the dying senescent BMSCs. Such post-death messages from the substantial amount of aging bone marrow stromal tissues may act as a vital regulator in aging-associated systemic immunosuppression and epithelial protumorigenesis. In malignant tissues, a dominant feature is the presence of polyploid giant cells (PGCs) which contain more than two sets of genomes through endopolyploidy and prevail in almost all senescent cells including MSCs (19–22). Based on the evidence of ghost messages produced by cell death including apoptotic cell death (23, 24), we hypothesize that post-death signals may arise from dying PGCs within senescent BMSCs, contributing to an aging-related factor associated with immune cell suppression and deterioration of breast epithelium. Using replicative senescent healthy female BMSCs, we examine the function of PGCs in the juvenile and advanced phases of senescence. We discover the formation of post-death vesicles termed Senescent PGC-Spiking Bodies (SPSBs) originating from dying senescent PGCs, which accelerate the risk and progression of breast cancer by promoting macrophage aging with facilitated cancer-prone epithelial transition.

## RESULTS

### Biphasic PGCs in the course of fBMSCs Senescence

To explore the possibility that aging bone marrow stromal cells may produce signals that predispose individuals to breast cancer, we undertook an in-depth analysis of PGCs throughout senescent course of BMSCs from a cohort of healthy adult female bone marrow donors (age 19-52, n = 8; Supplementary Fig. S1A). The fBMSCs were identified by BMSCs and non-BMSC biomarkers (Supplementary Fig. 1B,C), and went through the replicative senescence following the established in vitro expansion (25, 26, 27). Based on PGC morphologic features and karyotypic alternations across the fBMSC donors (Fig. 1A-C, Supplementary Fig. S2A-C), the status of fBMSC senescent progression was divided into juvenile (phase I; passage 1-7), senescing (phase II; passage 8-12) and advanced-senescent (phase III; passage >13). A baseline range of PGC population (5-10%) was detected in the juvenile phase fBMSCs across the donors (Fig. 1C and Supplementary Fig. S2B). As fBMSCs proceeded through the senescing to the advanced senescence, the senescent SA-β-gal^+^/p16^+^ PGCs were gradually increased to as high as ∼40% with remarkably reduced proliferative Ki67 and DNA repair activity (Fig. 1D-F, Supplementary Fig. S2D). Surprisingly, the PGCs in the juvenile phase fBMSCs exhibited the proliferative and DNA repair capacity measured by Ki67 and γH2AX/53BP1 foci formation after radiation (Fig. 1F, Supplementary Fig. S3). In addition, contrasted with the PGCs in the senescing and advanced-senescent fBMSCs, PGCs in the juvenile phase showed a notable tendency of contacting with neighboring cells for clonal initiation and expansion (PGCs, red arrows, proliferative diploid cells, blue arrows; Fig. 1G, Supplementary Fig. S4A,B). The PGCs were found to be preferably located at the edges of the growing colony (Fig. 1G, Supplementary Fig. S4C) and the percentage of PGCs/colony decreased with the predominant proliferation of diploid fBMSCs as colonies underwent expansion (Fig. 1H). The PGCs isolated from fBMSCs colonies were able to boost fBMSC proliferation (Fig. 1I) and the PGCs/colony ratio was dose-dependently increased in radiation-surviving colonies (Supplementary Fig. S4D-F). In contrast, such proliferation-stimulating PGCs were reduced in the senescing phase and absent in the advanced-senescent fBMSCs (phases I, II, Fig. 1, Supplementary Fig. S4B). In consistence, a dandelion seeding-like cell budding phenomenon was detected in female mouse BMSCs (fmBMSCs) isolated from young but not old mice (Supplementary Fig. S5). Taken together, these findings indicate that the PGCs formed in the juvenile phase fBMSCs can enhance clonal expansion via boosting neighboring diploid cell proliferation (Fig. 1J), a potential vital function for maintaining BMSC homeostasis and BMSC-mediated tissue regeneration.

**Fig. 1.**
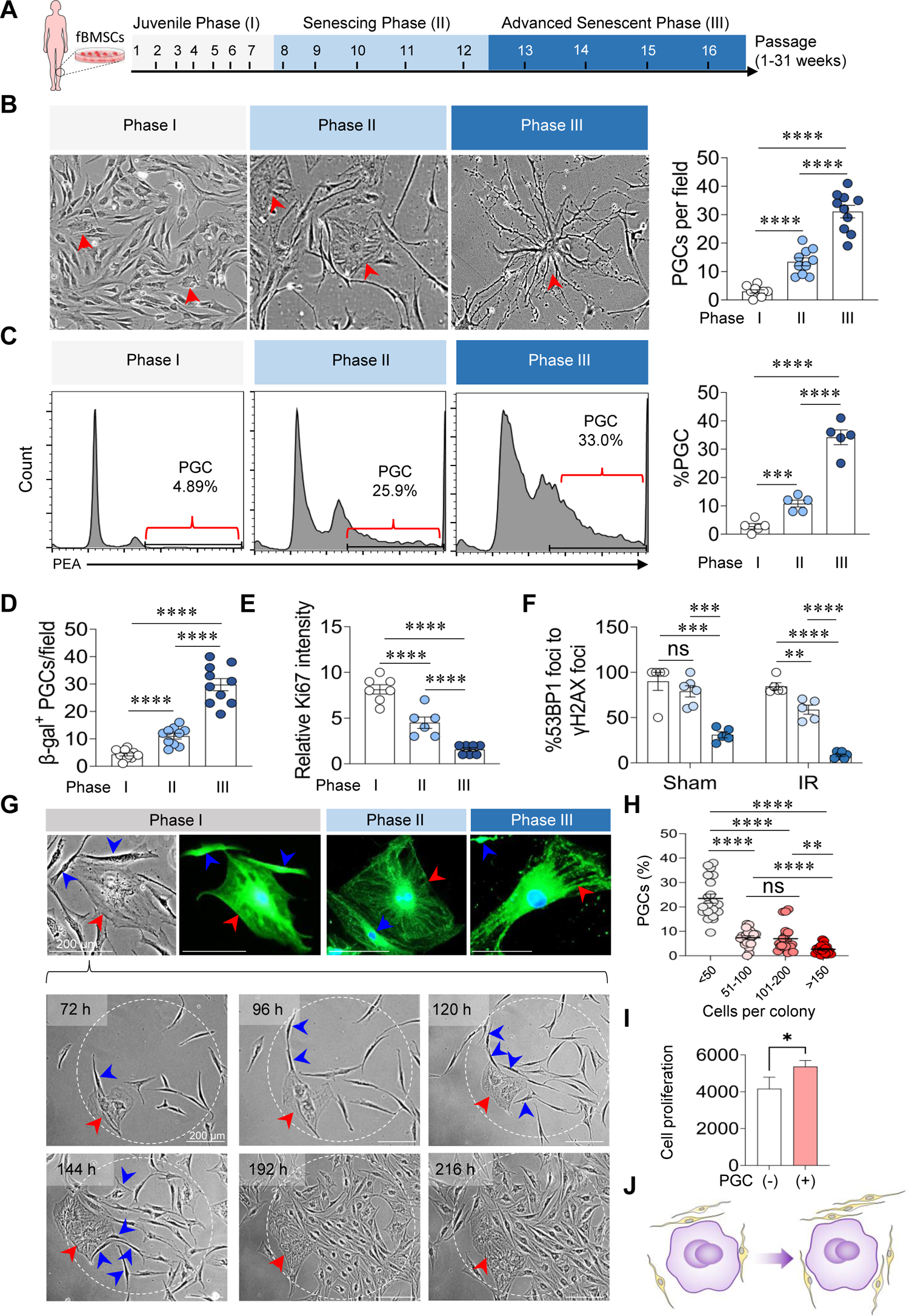
Biphasic PGCs in fBMSCs juvenile-to-senescent progression. **A**, Schematic time course of fBMSC senescence monitored in juvenile (phase I), senescing (phase II) and advanced senescent (phase III) phases. **B**, Morphologic features of fBMSCs and PGCs in the indicated senescent phases (left) and PGCs calculated per randomly selected areas (right, n = 10). **C,** PGCs measured by karyotyping (n = 5). PGCs in the indicated senescent phases of fBMSCs were tested by senescent biomarker SA-β-galactosidase (**D**), proliferative biomarker Ki67 (**E**) and DNA repair activity by 53BP1 foci (**F**); in D-F, n = 5-10. **G**, PGCs (red arrow) with (phases I and II) or without (phase III) connection with neighboring diploid fBMSC cells (blue arrows). Lower panel: time-lapse images of a PGC (red arrow) boosting clonal proliferation of fBMSCs (blue arrow). **H**, Percentage of PGCs per colony calculated at different stages of fBMSC clonal development (n = 20). **I**, Growth of GFP-fBMSCs measured by flow cytometry following co-culture with PGCs isolated from fBMSC colonies for 72 hours; co-culture of GFP-fBMSCs with fBMSCs as the negative control (n = 3). **J**, Schematics of PGC boosting neighboring fBMSCs in clonal proliferation.

### Post-death Vesicles Generated from Dying PGCs in Advanced-senescent fBMSCs

To evaluate potential ghost signals from dying-senescent PGCs in the advanced-senescent fBMSCs we monitored the PGCs as fBMSCs progressed through the senescing to the advanced-senescent phase which displayed the distinctive senescence-related cellular and mitochondrial degenerations (refs 22, 28, Fig. 2A, B). The dying PGCs demonstrate a four-step disassembly process observed in live conditions using bright field microscopy or dark field with IF of MitoTracker Red. Step 1, mitochondrial fusion and elongation, Step 2, formation of mitochondria-containing spikings, Step 3, spikings are beaded into membraned vesicles (termed Senescent PGCs Spiking Bodies, SPSBs), and finally, in Step 4, more than 50% of the spikings converted into SPSBs (Fig. 2B, Supplementary Video S1 and S2). The spiking-derived SPSBs were then characterized by potential biomarkers with IF of an array of lysosomal and mitophagy biomarkers, showing co-staining of p62 and RAB7 (Fig. 2C, top panel, yellow arrows) both involving mitophagosome traffic and degradation. In addition, SPSBs were remarkedly enriched with the tetraspanin protein CD63 co-stained with p62 and MitoTrackerRed (Fig. 2C, middle and lower panels, yellow arrows, Supplementary Fig. S6A,B). In addition to that CD63 plays a pivotal role in the orchestration of the endosomal cargo and synthesis of EVs, SPSBs also express a cluster of lysosomal and mitochondrial biomarkers (summarized in Supplementary Fig. S6C). The SPSBs were isolated from the advanced-senescent fBMSCs enriched with the spiking-PGCs by a series of conditional trypsinization and centrifugation without detaching the PGCs (Method, Supplementary Fig. S6D). SPSBs were identified to be membraned vesicles containing mitochondrial structures by transmission electron microscopy (TEM) with Cryo-EM and negative staining (Fig. 2D, yellow arrows, Supplementary Fig. S6E) with an average volume dimension of 200-400 nm calculated by NTA and TEM (Fig. 2E). mtDNA was confirmed through PCR and sequencing identification (Fig. 2F, Supplementary Fig. S6F,G). These results suggest that SPSBs are a unique post-death vesicle from the dying PGCs in the advanced-senescent fBMSCs. SPSBs could be a regulator contributing to bone marrow aging-related pathological status including immunosuppression and BC risk.

**Fig 2.**
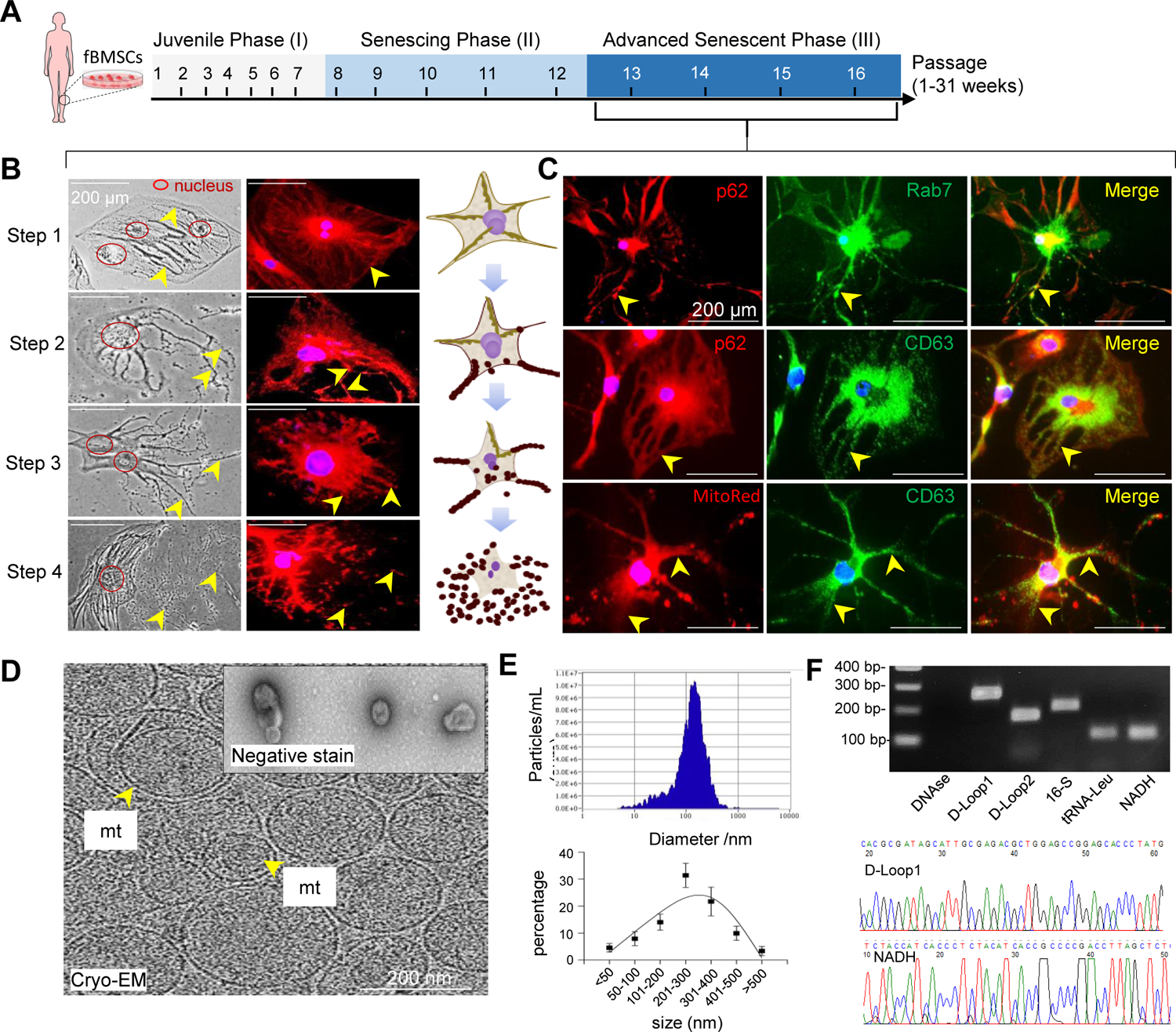
SPSB generation by dying PGCs in advanced-senescent fBMSCs. **A**, Schematic time course of fBMSC senescence monitored in juvenile (phase I), senescing (phase II) and advanced senescent (phase III) phases. **B,** Representative dynamics in SPSB generation (yellow arrows) by dying-senescent PGCs through 4-step mitochondria-containing spike formation and disassembling (bright field and MitoTrackerRed). **C,** Immunofluorescent (IF) images of SPSBs tested by lysosomal and mitophagy biomarkers with different combinations: CD63 and MitoTrackerRed (top), CD63 and p62 (middle), and Rab7 and p62 (bottom); nucleus stained by DAPi, blue. **D,** Cryo-electron microscopic images of SPSBs (yellow arrow indicating the mitochondria-like structure parceled in SPSBs; insert, negative staining). **E,** SPSBs volume distribution measured by NTA (top) and by TEM (bottom). Percentage and size distribution of SPSBs. **F,** PCR of SPSB mtDNA with indicated mtDNA fragments (left) with identified mtDNA sequences (right). * *p*< 0.05, ** *p*< 0.01, *** *p*< 0.001, **** *p*< 0.0001, ns, *p*> 0.05, data are shown as mean ± SEM.

### SPSBs Accelerate Macrophage Aging and Reduce Phagocytosis with Protumor Polarization

Free mtDNA released from damaged cells has been shown to dysregulate immune cells including macrophage suppression which plays a critical role in BC progression (29). We postulated that macrophages constitute a primary target for SPSBs due to the potential immunosurveillance role of macrophages in clearing aging-BMSC-derived SPSBs. Indeed, SPSBs generated from dying GFP-PGCs can be rapidly endocytosed by macrophages differentiated from THP-1 monocytes by phorbol myristate acetate (PMA) (Fig 3A, Supplementary Fig. S7A). THP-1 cells differentiated with PMA+SPSBs explicated a significant proliferation enhancement (Fig. 3B,C) and accelerated maturation indicated by CD68^+^ population CD68 IF (red arrows, Fig. 3D,E, Supplementary Fig. S7B) which could not be further enhanced by extending the time of SPSB+PMA treatment (Fig. 3F), suggesting a maximal level of macrophage maturation promoted by SPSBs. The SPSB+PMA maturated macrophages accelerated senescence measured by SA-β-Gal expression which was further enhanced by extending the time of SPSB exposure (Fig. 3G,H, Supplementary Fig. S7C,D). In addition, SPSB+PMA differentiated macrophages exhibited impaired phagocytic clearance on MCF-7 cells (Fig. 3I) and resistance to M1 polarization induced by LPS and IFNγ, while concurrently accelerated M2 polarization induced by IL6 (Fig. 3J, Supplementary Fig. S7E,F). In consistence, IL-6, IL-1β, and TNFα which recruit anti-tumor immune cells and induce cytotoxic immune responses remained unchanged in the SPSB+PMA-induced macrophages (Fig. 3K) whereas IL-10 that suppresses immune responses, CCL22 that attracts Treg cells, and PPARγ that enhances M2 macrophages, were all enhanced in the SPSB+PMA-induced macrophages (Fig. 3L). Thus, agreed with mtDNA-inhibited immune regulation (30), the SPSBs carrying the mtDNA and other deteriorative signals can promote macrophage aging, a vital factor in breast cancer initiation and progression.

**Fig 3.**
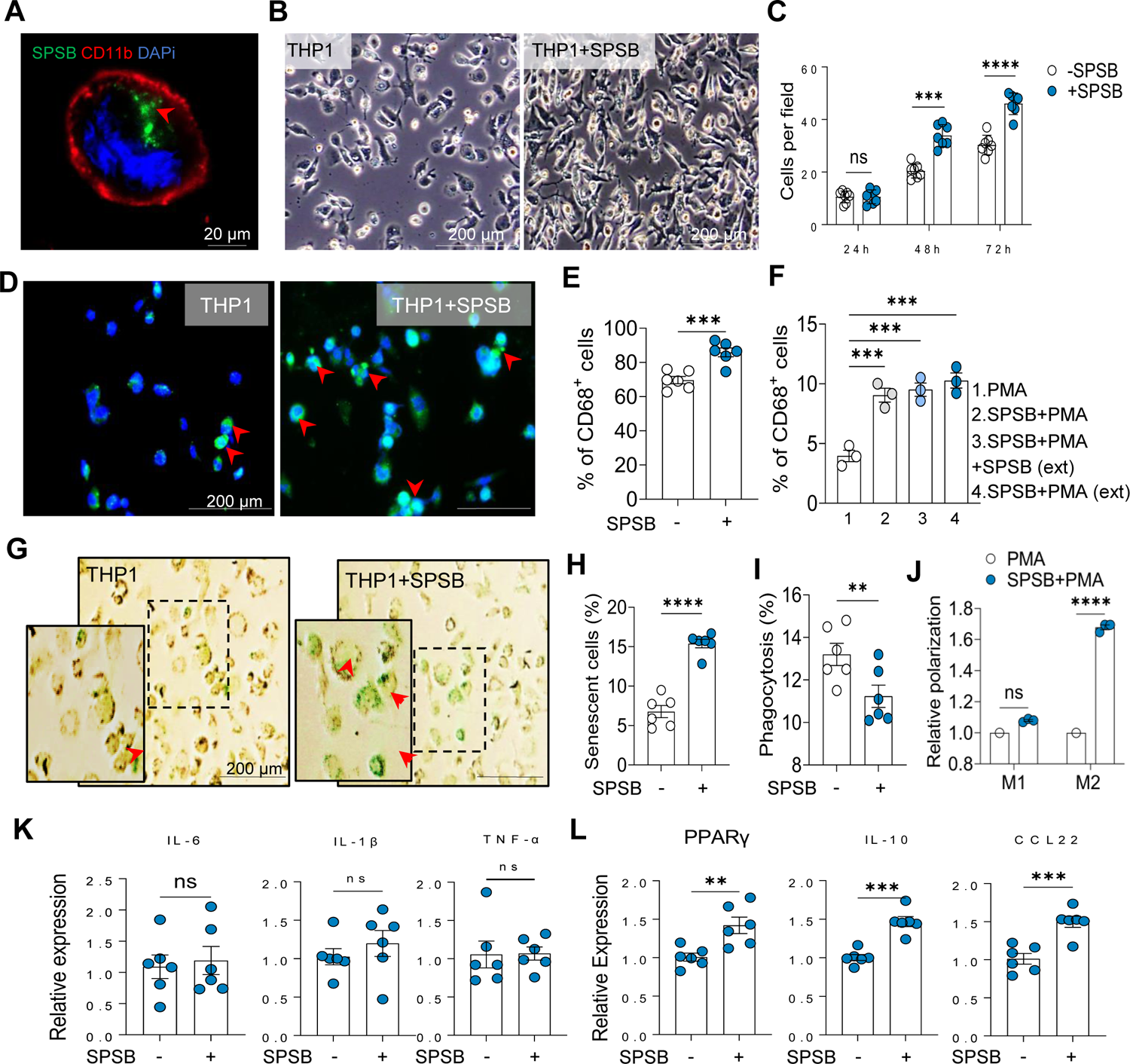
SPSBs accelerate macrophage aging and impair phagocytosis in BC cells. **A**, Representative IF image of macrophage endocytosed GFP-SPSBs generated from GFP-PGCs in advanced-senescent fBMSCs. The image illustrates the monocyte/macrophage biomarker CD11b (in red) on the cell membrane and the endocytosed SPSBs in the cytoplasm (in green) with nuclear DNA stained by DAPI (in blue). Images (**B**) and quantitation (**C**) depicting **e**nhanced THP-1 cell proliferation coculture with SPSBs (1250/cell) and PMA (40 nM) for 72 hours; n = 7. IF (**D**) and flow cytometry (**E**) analysis of CD68^+^ cells in THP-1 cells differentiated by PMA, SPSB+PMA, or extended time (72 hours) of SPSB+PMA with or without SPSB, n = 3. **F,** Flow cytometry analysis of CD68+ THP-1 cells treated with different combination of SPSBs and PMA (n = 3). **G**. Representative images of SA-β-gal positive THP-1 cells following PMA treatment with or without SPSBs. **H,** Quantification of senescent THP-1 cells by ImageJ, n = 6. **I,** Flow cytometry measured phagocytosis of MCF-7 cells by macrophages pretreated with PMA in the presence or absence of SPSBs (n = 6). **J,** M1 or M2 polarization of THP-1 cells with or without SPSBs pretreatment followed by PMA (40 nM, 72 hours) incubation. M1 polarization with IL-6 (20 ng/ml 48 hours) or M2 polarization with IL-4 (20 ng/ml) and IL-13 (20 ng/ml) for 24 hour). n = 6. **K,** M1 cytokine mRNA levels expressed by THP-1 cells treated with PMA (100 nM) + SPSBs (1.5 µg/ml) for 72 hours followed by LPS (100 ng/ml) and IFNγ (20 ng/ml) induction. **l,** M2 cytokine mRNA levels expressed by THP-1 cells treated with PMA (100 nM) + SPSBs (1.5 µg/ml) for 72 hours, followed by IL-4 (20 ng/ml) and IL-13 (20 mg/ml) induction. ** *p*< 0.01, *** *p*< 0.001, **** *p*< 0.0001, ns, *p*> 0.05, data are shown as mean ± SEM.

### SPSBs Carried Mitochondrial Elements Are Enriched in BC and Associated with Poor Prognosis

mtDNA can boost tumor invasion (31, 32) and recently, mtDNA has been found to induce SASP in radiation-induced senescence which is blocked for enhancing mouse lifespan (31, 32). With SPSB-mediated macrophage senescence which plays a critical role in aging-associated BC risk and progression, we wondered, beyond the mtDNA, SPSB cargo may contain additional cancer-promoting signals. Indeed, SPSB proteomics versus control of juvenile fBMSC-EVs that induce tissue repair and regeneration (33), revealed a cluster of mitochondrial OXPHOS elements that are well-defined to induce redox imbalance in aging organs and BC tumorigenesis(34). A group of key elements, AIFM1, NDUFS1, NDUFAB1, and PPA2 within a large group of components of the mitochondrial respiration chain with categorized functions and biological processes (Fig. 4A, Supplementary Fig. S8A,B) were found to be significantly elevated in BC compared to surrounding normal tissues (Fig. 4B), and such tendency was also evidenced in tumor tissue of patients with age >= 55 years old versus the control group of patients with age <54 year old (Fig. 4C). Furthermore, enhanced expression of each OXPHOS gene was associated with a worse clinical outcome in BC patients (Fig. 4D). These results demonstrate that SPSB-carried mtDNA and the redox-inducing OXPHOS elements, together with SPSB-mediated macrophage aging, could promote breast cancer risk via transforming breast epithelial cells.

**Fig. 4.**
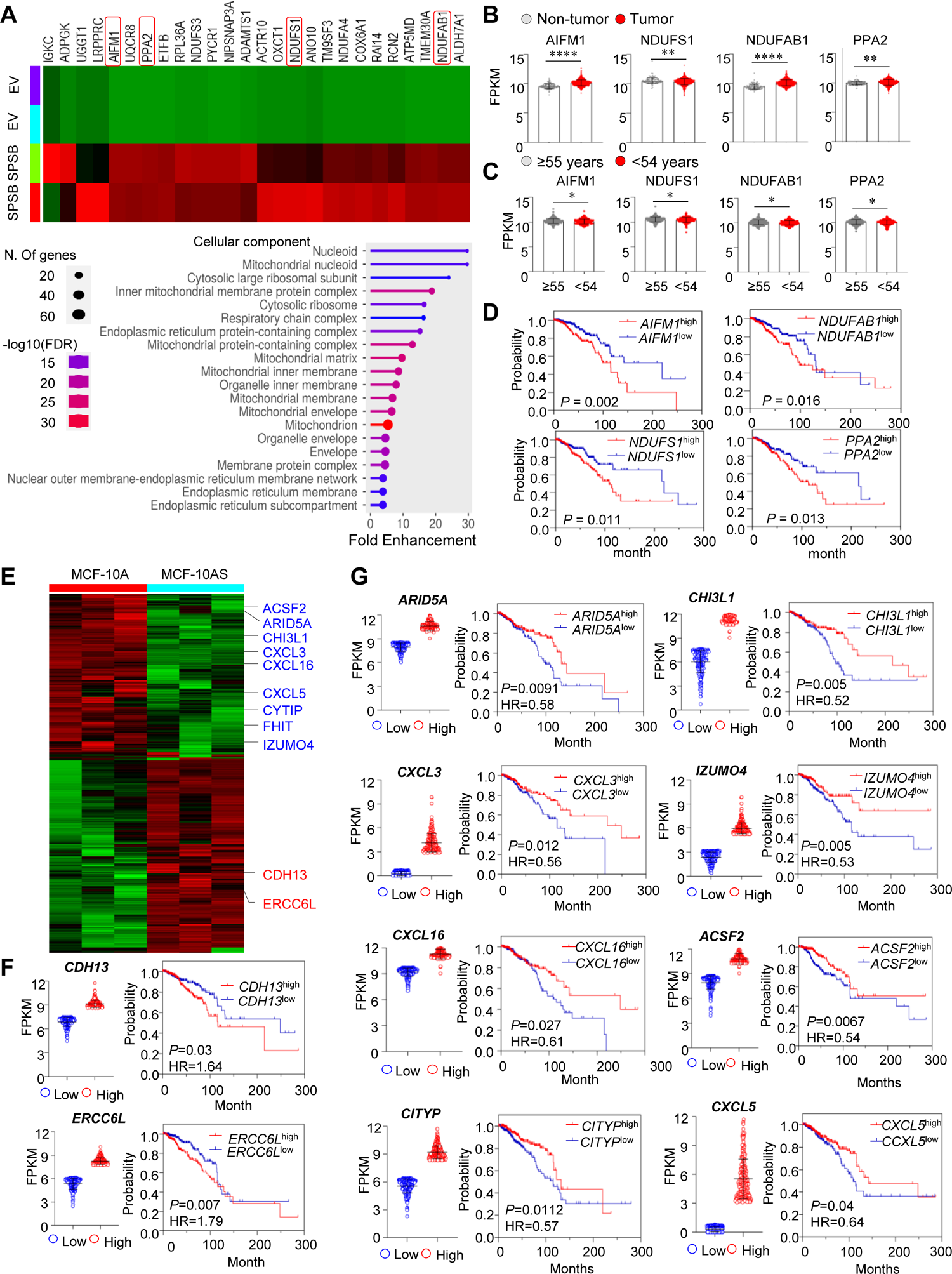
Mitochondrial respiration elements in SPSB cargo are elevated in BC and connected with elder patient prognosis; SPSB-induced BC oncogenic pattern is associated BC poor prognosis. **A,** SPSB cargo proteomics with the control senescing phase II fBMSC-EVs revealing an enriched cluster of mitochondrial OXPHOS elements in SPSBs; lower panel, fold enhancement of the cellular component of SPSB cargo proteins compared to hBMSCs-EVs. **B,** Expression of OXPHOS elements in BC (n = 1097) versus non-tumor surrounding tissues (n = 114). **C,** Expression of OXPHOS elements in young (≤54 years old, n = 338) and elder (≥55 years old, n = 480) BC patients. **D,** Enhanced mitochondrial OXPHOS elements (AIFM1, NDUFS1, NDUFAB1, and PPA2) connect with BC patient poor prognosis by comparing the top (n = 272) and bottom (n = 272) quartiles of the indicated subgroups in TCGA database. **E,** Transcriptomic profile of SPSB-promoted MCF-10AS cells versus the counterpart control MCF-10A cells revealing the SPSBs-upregulated oncogenes (*CDH13*^high^ and *ERCC6L* red) and the down-regulated tumor suppressor genes (blue). **F,** SPSB-upregulated oncogenes associated with poor overall survival (OS) of BC patients by comparing the top (n = 275) and bottom (n = 275) quartiles of *CDH13*^high^ and *CDH13*^low^ subgroups or *ERCC6L*^high^ and *ERCC6L*^low^ subgroups in TCGA database. **G,** SPSB-down-regulated tumor suppressor genes, *ARID5A*, *CHIL31, CXCL3, CXCL16*, and *IZUMO4* connected with the OS by comparing the top (n = 275) and bottom (n = 275) quartiles of the indicated subgroups in TCGA database. The same cut-off of segregation between the highest and lowest expression was used as above. * *p*< 0.05, ** *p*< 0.01, **** *p*< 0.0001, ns, *p*> 0.05, data are shown as mean ± SEM.

### SPSBs Induce Breast Epithelial Cell Protumorigenic Transition

Agreed with the well-established redox-induced BC initiation and progression(35), transcriptomic analysis of SPSB-incorporated human normal breast epithelial MCF-7 cells (MCF-10AS) revealed a specific oncogenic pattern including two upregulated tumor-promoting genes (*CDH13* and *ERCC6L*) and downregulate tumour suppressor genes (*ACSF2*, *ARID5A*, *CHI3L1*, *CXCL16*, *CXCL3*, *CXCL5*, *CYTIP*, *FHIT* and *IZUMO4,* Fig. 4E). BC database analysis revelated that prognosis of BC patients is associated with the SPSB-upregulated oncogenes (Fig. 4F) and SPSB-down regulated tumor suppressor genes (Fig. 4G). With the above SPSB-mediated macrophage aging and the pro-BC redox-inducing elements, and the reported interdependent macrophage dysfunction and BC progression (29, 36), we further investigated the SPSB-mediated epithelial tumorigenic potential. Firstly, a higher frequency of MCF-10A clonal expansion attached to the PGCs rather than the fBMSCs was detected by coculturing MCF-10A cells with the senescing fBMSCs (Fig. 5A, Supplementary Fig. S8C) and SPSBs can directly boost MCF-10A cell proliferation (Fig. 5B). Like SPSBs targeting macrophages, SPSBs were promptly endocytosed by MCF-10A cells, as visualized by GFP-labeled SPSBs located in the cytoplasm of RFP-labeled MCF-10A cells (yellow arrows, Fig. 5C). Consistent with the pro-BC transcriptomic pattern (Fig. 4E), MCF-10AS cells showed protumorigenic cell morphology (red arrows, Fig. 5D) with enhanced clonogenicity (Fig. 5E, Supplementary Fig. S8D). A cluster of genes for cell migration and cancer metastasis including MMP28 (epilysin) promoting cancer invasion, NECTIN3 in cell migration, DCN regulating collagen fibril organization and migration, FYN enhancing cancer metastasis, Contactin-1/F3 in neuronal migration, and SNAI2, a transcription factor regulating EMT and tumor invasion, was identified in MCF-10AS transcriptome (Fig. 5F,G). In consistency, MCF-10AS cells accelerated wound-healing ability (Fig. 5H,I), Matrigel penetration (Fig. 5J,K, Supplementary Fig. S8E), and depolarized 3-D acini structures with epical-basal disorganization (Fig. 5L,M, Supplementary Fig. S8F). Despite the absence of fully transformed tumors, MCF-10AS cells generated precancerous acini compared to MCF-10A cells inoculated in the opposite mammary pat (Fig. 5N) showing the precancerous feature in the gland structures, deviating from the regular hollow lumen glands formed by control MCF-10A cells (Fig. 5O). The GFP-expressing acini with the protumourigenic features were confirmed by fluorescence IHC of Laminin V for basal membrane (red) and nucleus stained with DAPi (blue) and quantified in both mammary glands (Fig. 5P). The results indicate that SPSBs are not solely capable of initiating tumor, they can create a protumorigenic epithelial structure that, when accompanied by aging macrophage or other aging-related factor, can facilitate a malignant transformation.

**Fig. 5.**
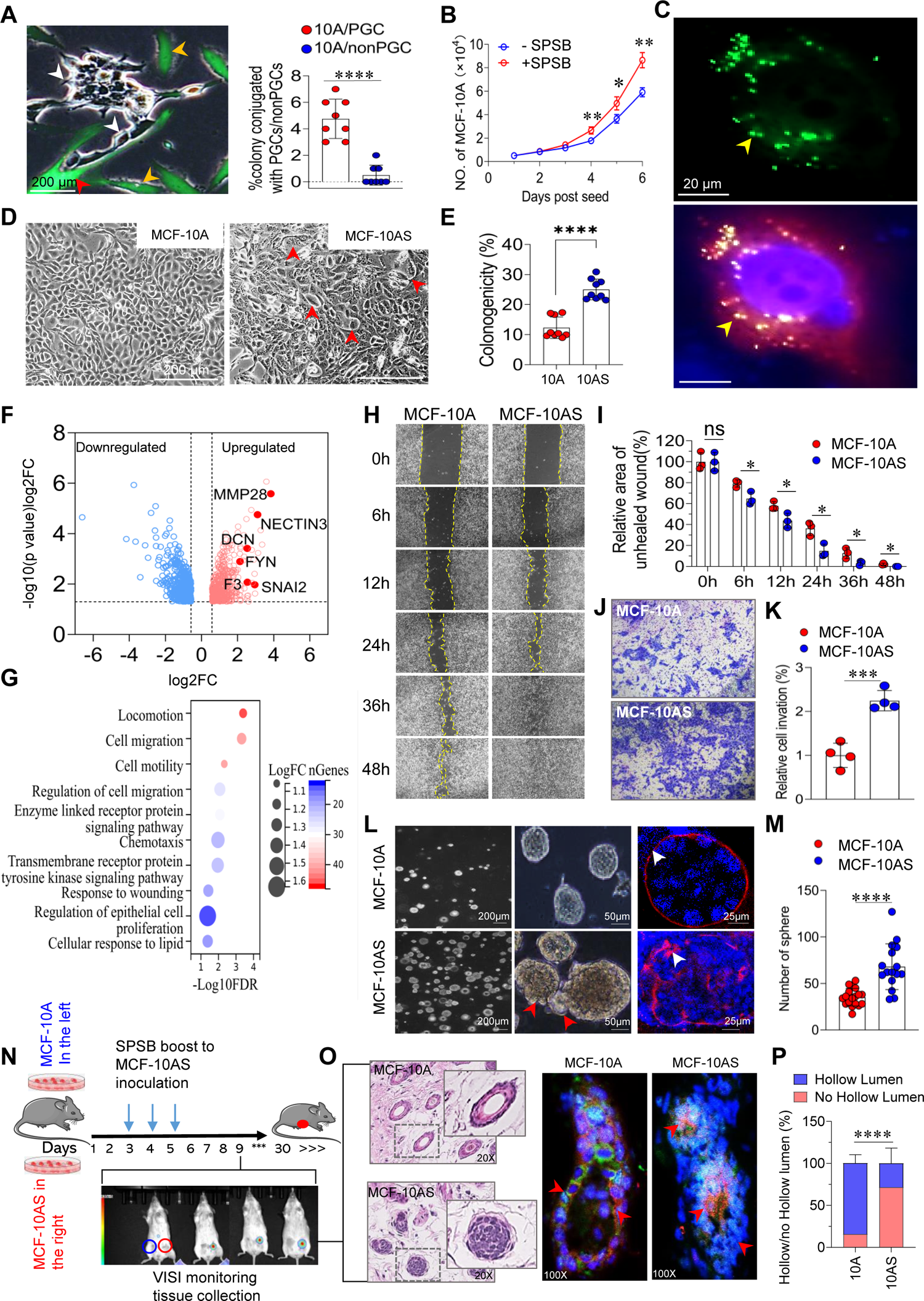
SPSBs promote breast epithelial cell protumorigenic transition. **A,** Images of a living growing MCF-10A colony attached to a GFP-PGC in phase II fBMSCs; red arrow, GFP-PGC, white arrow, clonal MCF-10A cells, gold arrow, fBMSCs. **B,** Quantified frequency of interactions of MCF10A colony/PGCs and MCF10A colony/fBMSCs (n = 8). **C,** MCF-10A cell proliferation and clonogenicity measured with or without coculture of SPSBs (n = 3). **D,** Endocytosed SPSBs (green, left) in an RFP-MCF-10AS cell with DAPi stained nucleus (blue, right). **E,** Disturbed cellular polarity and enlarged cells in long-term SPSB-promoted MCF-10A (MCF-10AS) cells compared to MCF-10A cells with similar passaging but without SPSB incorporation. **F,** Volcano plot based on fold change and *P* value of all transcripts in MCF-10AS versus MCF-10A cells. **G,** Pathways of cell migration and motility enhanced in MCF-10 and MCF-10AS cells. **H,I,** Images and quantification of wound healing rate in MCF-10A and MCF-10AS cells (n = 3). **J,K,** Images and quantification of Matrigel invasion of MCF-10AS and MCF-10A cells (n = 4). **L,M,** Representative images of the bright field of mammosphere and IF of Laminin V for basal membrane (red) and nucleus stained with DAPi (blue) and quantification of mammospheres formed by MCF-10A (n = 18) and MCF-10AS (n = 16) cells. Schematic of in vivo breast plugs acini analysis (**N**) and representative images of HE and IF Laminin V (red) with nucleus stained by DAPi (blue) (**O**) and quantitation (**P**) of hollow and non-hollow lumen acini generated with plugs of GFP-MCF-10A and GFP-MCF-10AS cells in mouse mammary fat pat (n = 8). * *p*< 0.05, ** *p*< 0.01, **** *p*< 0.0001, ns, *p*> 0.05, data are shown as mean ± SEM.

### PGCs formed in the SPSBs-transformed Cells Resemble the Polyploid Giant Cancer Cells

Polyploid giant cancer cells (PGCCs) are believed to be the transformed stem cells driving aggressive growth in solid tumor including BC (37, 38). SASP is shown to favorably transform stem cells and CD63 is identified in cancer stem cells (39), which together with our finding that CD63 is a predominant biomarker of SPSBs, promoted us to test if SPSB-promoted protumorigenesis is associated with PGCC formation. Indeed, karyotyping revealed a time-dependent increase of PGC population ranging from 16.9% to 22.3% in MCF-10AS cells compared to the basal 4.50% PGCs in MCF-10A cells with sham-SPSB culture condition; similarly PGC enhancement from 6.56% in MCF-10A cells to 17.3% PGCCs in MCF-10A cells transformed by overexpressing Her2, 11.3%, and 17.2% in MCF-7 and MDA-MB-231 cells (Supplementary Fig. S9A). The PGCs in MCF-10AS cells exhibit preferably location at the growing edges of wound healing (Fig. 6A,B) and early clonal expansion, mirroring the PGCCs in the clonal expansion of BC cells (Fig. 6C, Supplementary Fig. S9B,C). Analogous to PGC in juvenile fBMSC clonal expansion, the ratio of PGCs/colony and PGCCs/colony in MCF-10AS and BC cells decreased with clonal expansion due to diploid cell proliferation shown by karyotyping of pooled nature colonies (Fig. 6D) and calculated PGCs/colony and PGCCs/colony ratio (Supplementary Fig. S9D). Thus, formation of polyploidy seems to be required for cells under the pressure of fast proliferation. We then explored the possibility that the PGCs in MCF-10AS cells and PGCCs in BC could result from SPSB-transformed stem cells. Since CD63 is the dominant biomarker of SPSBs, we tested the CD63 expression in clonal BC cells and CD63^+^PGCs were detected in the clonal expansion of MCF-7 and MDA-MB-231 cells (Fig. 6E). Compared to the colonies of MCF-10A cells, numbers of CD63-expressing PGCs and CD63 expression levels were enhanced in the MCF-10AS colonies which were comparable to the PGCCs in the colonies formed by MCF-7 and MDA-MB-231 cells (Fig. 6F, Supplementary Fig. S10A-C). Furthermore, the CD63^+^PGCCs were identified in mouse spontaneous breast tumors induced by DMBA/MPA carcinogenic promotion as well as in the tumors with diagnosed BC subtypes (Fig. 6G-I). These results suggest that (summarized in Fig. 6J) PGCs formed in proliferative juvenile fBMSCs promote neighboring cell proliferation whereas advanced-senescent fBMSCs formed PGCs can produce SPSBs which carry deteriorating mtDNA and OXPHOS redox-inducing elements, induce macrophage aging and breast epithelial precancerous transition, two essential events in aging-associated breast cancer risk and progression.

**Fig. 6.**
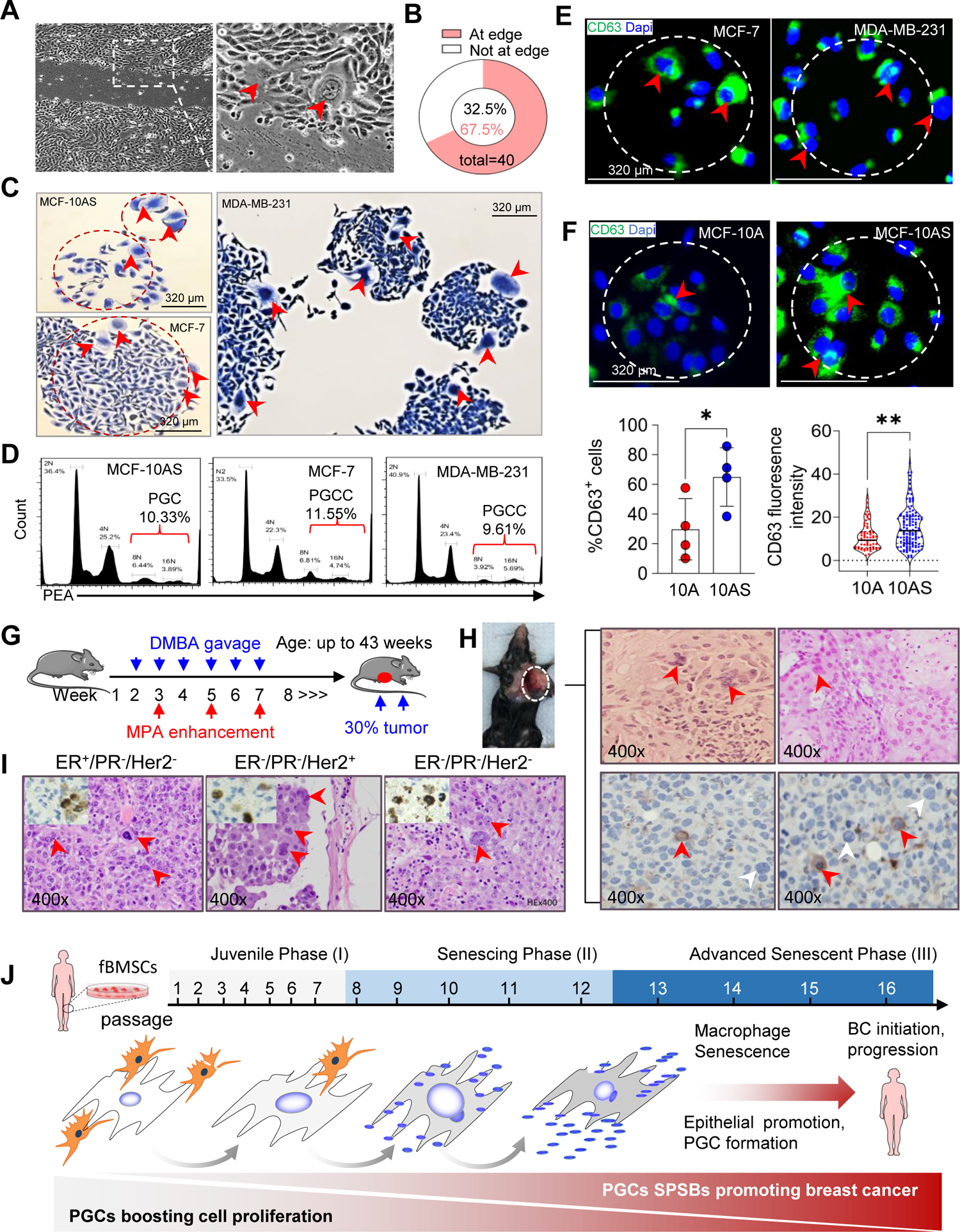
PGCs in clonogenic MCF-10AS bear resemblance to PGCCs in breast cancer. **A,B,** Living images and quantitation of PGCs (red arrow) calculated in the region within 50 cells and >50 cells from growing edges of wound healing MCF-10AS cells, n = 40. **C,** Representative PGCs and PGCCs (red arrow) in growing colonies of MCF-10AS and BC cells. **D,** Karyotyping of PGCs and PGCCs in colonies of C. **E,** Representative starting colonies with IF of CD63-PGCCs in BC cells (red arrow; CD63 = green, nucleus = blue). **F,** Representative IF and quantitation of CD63-PGCs in colonies of MCF-10A and MCF-10AS, n = 4 for measuring percentage of CD63-expressing cells; n = 40-60 for measuring CD63 fluorescence density. * *p*< 0.05, ** *p*< 0.01. **G,** Time scheme of DMBA/MPA induced mouse spontaneous breast tumor. **H,** Representative image of PGCCs in two mouse breast tumours identified by HE staining (top, red arrow) and IHC of CD63-PGCs (bottom). **I,** PGCCs by HE (red arrows) and CD63-expressing PGCCs (inserts) of human breast cancer with indicated subtypes. **J,** Schematic summary of breast cancer risk influenced by aging bone marrow stromal cells.

## DISCUSSION

Here we describe a previously unexplored post-cell death vesicle Senescent PGC Spiking Bodies (SPSBs) from dying PGCs in bone marrow mesenchymal stromal cells of healthy women undergoing replicative senescence. The SPSBs carrying mtDNA and OXPHOS redox elements are implicated in predisposing individuals to breast cancer by dual deteriorative macrophage senescence and breast epithelial cell protumorigenic transition. In line with the broader concept of “ghost messages” arising from acute cell death via apoptosis (23, 24), our findings unveil a breast cancer-promoting post-death message originating from dying-senescent bone marrow stromal cells. The bioactive elements encapsulated and shielded by SPSBs may offer the advantage of facilitating and maintaining distant communication for aging-related pathogenesis, thus representing a potentially effective target for attenuating age-associated degenerative conditions and reducing cancer risk (1, 2, 40). The degenerative signals carried by senescent PGCs, along with the proliferation-stimulating function of PGCs in the juvenile phase of fBMSCs, explicate the dynamics of PGC behavior influenced by the environments of host cells with proliferative or senescent status.

Contrasted to the well-characterized SASP and senescent cell-derived EVs secreted by metabolically active cells (17, 41), SPSBs are generated by metabolically inactive (Ki67-negative) PGCs and share biomarkers of mitophagy including CD62 and CD63. Unlike mitophagy occurring within the cytoplasm, SPSBs are directly shed from mitochondria-enriched spikings through a morphologically distinct 4-step process. In addition, the cargo contents of SPSBs also indicate a unique message, characterized by predominant redox-inducing elements. Such aging-BMSC induced messages may undergo specific enhancement during the terminal stage of chronic cell death in aging stromal tissues, different from the biological function induced by instant free mtDNA not protected by membraned vesicles (42) and released from acute cell death after stress-induced senescence and apoptosis (31, 32, 43). Thus, instead of acute proinflammatory response in the local tissue induced by SASP, cellular redox imbalance induced by the SPSB-targeted distant tissues may accelerate organ degenerative conditions, leading to immune cell senescence and cancer initiation and progression (44) which may be particularly critical in transforming stem cells in breast cancer progression (35). The concept of SPSB-distant signal delivery of mtDNA and redox-related OXPHOS elements concur with the report that circulating EVs of aged animals contain a higher level of ROS than the young animal (45). In the targeted immune or epithelial cells, SPSB-induced redox imbalance and the subsequent genomic and epigenomic regulation may induce different cellular response inducing the stress granules formed by condensing myriad proteins with mRNAs in the cytoplasm (46). However, despite the cellular stress response induced by SPSBs, it appears that this alone may not be adequate to induce malignant diseases. In in vivo studies, no fully transformed malignant tumors were identified with SPSB-promoted breast epithelial cells, although notable aggressive cell behavior and protumorigenic acini were observed. This suggests that post-death messages from aging bone stromal cells may not independently initiate malignant cell transformation but may instead synergize with other carcinogenic agents such as estrogen exposure in breast cancer risk (47).

A similar growth-stimulating feature was detected among the PGCs in juvenile proliferative fBMSC, SPSB-promoted breast epithelial MCF-10As cells and the well-observed PGCCs in breast cancer cells, although the complex genomic and transcriptomic identification are to be revealed. Both PGCs and PGCCs are specific cellular types containing more than two sets of genomes contained in multiple nuclei or a giant nucleus in a heterogenic cell population. Formation of PGCs is generally attributed to the genomic structural lability (37), and led to the “virgin birth” in cancer origination (48). The mechanism causing PGCs and PGCCs include endoreplication, mitotic failure and cell fusion which can be induced in cancer cells by genotoxic stresses including cancer radiotherapy (49). We found that PGCs within the proliferating fBMSC exhibit a cell-boosting capacity akin to the colony-initiating function observed in the clonogenic expansion of SPSB-promoted MCF-10AS cells and BC cells. Based on the similar roles of PGCs in normal and malignant tissues(37, 50), especially PGCCs in initiation of cancer stem cells (51), a functional syncytium of PGCs and PGCCs seems to present for clonal expansion in rapidly proliferating cells. This partially supported by the finding that the CD63 which is a prominent biomarker of SPSBs is expressed in the PGCCs in clonogenic MCF-10AS, BC cells, as well as spontaneous mouse breast tumor and clinic diagnosed BC. Although it is unclear why the CD63-carrying SPSBs can enhanced CD63 expression in the PGCCs, SPSBs may specifically target stem cells to form PGCCs as evidenced by stem cells preferentially targeted by SASP(52). The PGCCs via stem cell transformation could be resulted from integrated signaling from both SPSBs and SASP that induces chronic low-level inflammation, termed “inflammaging” determining the rate of stem cell transformation in BC initiation and organ aging (53, 54). The magnitude of amalgamated signals targeting stem cells from SASP in local tissue and SPSBs from systemic aging stromal tissue may represent the “natural selective force” underlying early-versus late-life antagonistic pleiotropy (55, 56). This phenomenon, as described by the pleiotropic genes being enhanced to adapt to the environment during the juvenile phase of life but declining with aging, may contribute to global immunosuppression and the propensity for cancer-prone epithelial deformation.

## METHODS

### Antibodies and Reagents

The staining for flow cytometry and immunofluorescence was performed at 4 °C in the presence of the indicated antibodies. The following antibodies were used in this study: FITC-conjugated anti-human IgG isotype (BD Bioscience, 555786), FITC-anti-CD14 (BD Bioscience, 555397), FITC-anti-CD34 (BD Bioscience, 555821), FITC-anti-CD44 (BD Bioscience, 555478), FITC-anti-CD105 (BD Bioscience, 561443), FITC-anti-CD68 (BioLegend 333805), PE-conjugated anti-human IgG isotype (BD Bioscience, 555787), PE-anti-CD45 (BD Bioscience, 555483), PE-anti-CD73 (BD Bioscience, 550257), APC-conjugated anti-human IgG isotype (BD Bioscience, 553932), APC-anti-CD90 (BD Bioscience, 559869), APC-Cy7-anti-CD11b (Biolegend, 101225), APC-Cy7-anti-CD206 (BioLegend, 321119), Alex488-anti-CD86 (BioLegend, 374203), Alex488-anti-CD80 (BioLegend, 375405), anti-53BP1 (Cell signaling, 4937), anti-phospho-Histone H2A.X (Ser139) (Millipore Sigma, 05-636), anti-Ki67 (Novus, B110-89717), anti-CD146 (Miltenyi Biotech, 130092849), anti-LamininV (Chemicon MAB19562), anti-CD11b (Bio-Rad, MCA551F), anti-α-Tubulin (Novus, NB100-690). The following secondary antibodies was used: Goat anti-Rabbit IgG-AF488 (Invitrogen, A11008), Goat anti-Rat IgG-AF647 (Invitrogen, A21247) and AF488-goat anti-mouse (A11017) from Invitrogen. Aexa-488-conjugated goat anti-rabbit IgG (A11034) and Goat anti-Rabbit IgG-Alexa-594 (Invitrogen, A11072), Rhodamine Red-conjugated goat anti-mouse antibody (Jacksonimmuno, 115-295-003), DAPI (Millipore Sigma, BD0015). Other reagents were also used: Poly-L-Lysine (Millipore Sigma, P4832); EasyPep Mini MS sample prep Kit (Thermoscientific, A40006). Fast SyBR Green Master Mix (Appliedbiosystem, 4385612), SuperScriptIII First-Strand kit (Life Technologies, 18080-51), Trizol (Life Technologies, 15596026), anti-CD63 (Invitrogen, 10628), anti-p62 (Sigma Millipore, P0067), anti-Rab7 (Cell Signaling, 95746), anti-CD9 (Sigma Millipore, CBL162), anti-Calnexin (Signaling, 2627T), CD81 (Sigma Millipore, MABf2061), anti-p21 (Proteintech Group Inc, 67362-1g), and anti-α-tubulin (Sigma Millipore, ABT170).

### fBMSCs

Human female bone marrow mesenchymal stromal cells (fBMSCs) were provided by the Institute for Regenerative Cures, University of California Davis, School of Medicine, from commercially available bone marrow (StemExpress) from 25 to 52-year-old healthy female bone marrow donors. Following established protocol of isolation and maintenance of BMSCs (57–59), fBMSCs were isolated from the mononuclear fraction using density gradient and adherence capacity. The cells were then cultured in low-glucose Dulbecco’s modified Eagle’s medium (DMEM) supplemented with 10% fetal calf serum (FCS) and identified by MSC biomarkers including CD73^+^, CD90^+^, CD105^+^, CD34^−^, CD45^−^ (refs 58, 59, Supplementary Fig. S1A-C). The fBMSCs from all donors were then maintained with regular passaging and monitored for PGCs up to 16 passages (31 weeks) which were characterized by juvenile, senescing, and advanced-senescent phases according to the morphological alterations, karyotyping, senescence biomarkers SA-β-galactosidase activity and p16, Ki67 proliferative index, and DNA repair activity.

### Human Breast Cancer Tissues

Human breast cancer pathological slides were provided by the UC Davis Comprehensive Cancer Center Biorepository. The acquisition of these samples was conducted in accordance with the regulations and guidelines set forth by the Institutional Review Board (IRB) under the protocol numbers 283665 and 218204. The UC Davis Comprehensive Cancer Center Support Grant (CCSG), granted by the National Cancer Institute (NCI P30CA093373), provided financial support for the biorepository. It is important to note that all patient information, except for the diagnostic results, has been anonymized to ensure privacy and confidentiality.

### Mice

All animal use and care protocols for isolation of mfBMSCs were approved by the Institutional Animal Use and Care Committee of the University of California Davis (IACUC 15315). Female Balb/c mice (JAX: 002019) and Female NRG mice (JAX:007799) were purchased from Jackson Laboratory All the animals were housed with HEPA-filtered ventilation and temperature-regulated cages kept at 21 or 29 °C. All of mice were maintained according to a standard guideline with free access to food and water.

### Other Cell Lines

Breast epithelial MCF-10A cells were purchased from ATCC and SPSB-promoted MCF-10AS cells were generated from PI’s Lab. Both cell lines were maintained in DMEM medium (Corning, 15-017-CV) with 5% horse serum, 20 ng/ml epidermal growth factor, 100 ng/ml cholera toxin (VWR, 80055-160), 0.5 mg/ml hydrocortisone (VWR, AAA16292-03), 10 mg/ml insulin (Sigma, I9278) and 1% penicillin/streptomycin (Corning, 30-002-CI). Breast cancer cell lines including MCF-7, MDA-MB-231 and 4T1, were maintained in DMEM medium supplemented with 10% FBS and 1% P/S. Human monocyte THP1 was purchased from ATCC and maintained in RPMI-1640 medium supplemented with 10% FBS, 1% P/S, 10 mM sodium pyruvate, 15 mM HEPES, 4.5 g/L glucose and 0.05 mM β-mercaptoethanol.

### PGCs Karyotyping

Cell suspensions of fBMSCs in phase I (juvenile, passage 2-7), phase II (sensing, passage 8-12), and phase III (advanced senescing) as well as cell suspension from SPSB-promoted MCF-10AS cells and pooled colonies from breast cancer cells were centrifuged and washed 3 times with PBS followed by resuspending the cells in PBS and fixed in 70% cold ethanol. The cells were then resuspended in 1 ml staining solution containing 0.2 mg/ml RNase A, 0.1% Triton-100, 0.1mg/ml BSA, and 0.05mg/ml Propidium iodide (PI). Cells were maintained in the dark at 37°C for 30 minutes prior to FACS analysis to calculate the percentage of PGCs gated with more than 4N genomic copies.

### Biomarkers of Senescent PGCs and Macrophages

The senescent status of PGCs in the progressing senescent phase of fBMSCs and SPSB-mediated senescent macrophages were recaptured following the established protocol (15, 60, 61). The senescence assay was conducted using Senescence β-Galactosidase Staining Kit (Cell Signaling, 9860). Briefly, cells were grown in 6-well plates and incubated with 1 ml of fixing solution for 15 minutes. After rinsing with PBS twice, 1 ml of β-Galactosidase staining solution was added and incubated at 37°C without CO2 overnight. SA-β-gal positive cells were visualized using Carl ZEISS microscopy GmbH. The assessment was performed in 10 randomly selected fields (20× magnification) for each individual sample.

### Fluorescence Immunocytochemistry

Cells were cultured on cover slides coated with a 0.01% Poly-L-Lysine solution and then fixed in 2% paraformaldehyde for 20 minutes. After washing and permeabilization with 0.5% Triton X-100 for 10 minutes on ice, the cells were treated with the staining enhancer Image-iT™ FX Signal Enhancer (Invitrogen, 136933) in a humid chamber at 37°C for 30 minutes. They were subsequently blocked with a solution containing 2% goat serum, 2% FBS, 1% BSA, 0.1% Triton X-100, and 0.05% Tween-20 for 1 hourat room temperature. Dual staining was performed by incubating the cells with primary antibodies at 4°C overnight. The cells were further exposed to corresponding fluorescent secondary antibodies. Nuclei were counterstained with DAPI. Cells were then visualized using Carl ZEISS microscopy GmbH. The fluorescence intensity was quantified using ImageJ.

### PGCs DNA Damage Foci Analysis

Juvenile, senescing and advanced senescent fBMSCs grown on coverslips in 60 mm cell culture dishes were irradiated at room temperature with 2 Gy X-ray (dose rate = 0.028 Gy/min; Hewlett Packard, McMinnville, OR, USA). Sham-irradiated coverslips were included as the controls. Cells were fixed 30 minutes post irradiation with 2% paraformaldehyde for 10 minutes followed by fluorescence immunocytochemistry staining for non-homologous DNA end joining (NHEJ) with incubation with the primary antibodies anti-53BP1 (Cell signaling, 4937) and anti-γH2AX (Cell signaling, 9718) overnight at 4°C. Foci images were acquired using a Zeiss LSM710 confocal microscope system with a 63× objective. Data analysis was carried out using ImageJ software. The number of repaired and unrepaired foci per nucleus was determined in 10-15 PGCs at different senescent phases.

### Time-lapse Cameras Recording PGCs

Conventional and confocal fluorescent microscopes were applied to monitor the living morphological changes of PGCs with or without GFP or RFP labeling for PGC-mediated neighboring cell proliferation and clonal expansion. High-throughput live-cell imaging system Incucyte S3 was added to monitor the PGCs in the advanced-senescent fBMSCs to capture SPSB generation from the dying-senescent PGCs. For the time-lapsing recording, different numbers of fBMSCs in senescing phase II or advanced senescent phase III were seeded in a 12-well plate and time-lap recording of 40-60 cells every 30 minutes for 2-4 days. The dynamic of dying-senescent PGCs and beading spikings were investigated with the records generated by Incucyte® S3.

### Clonogenic Assay

Cell clonogenicity and radiation clonogenic surviving of fBMSCs, SPSB-promoted MCF10AS and breast cancer cells were tested using our established protocols (62, 63). For measuring the clonogenicity of fBMSCs at different senescent phages, varied cell numbers of juvenile phase fBMSCs were seeded with different cell numbers and allowed to attach for 24 hours before being irradiated with X-rays at varying doses with the control cells unirradiated as the plating efficiency. Subsequently, the cells were incubated for 24-36 days for clonal development. The resulting colonies were stained using a 0.25% Coomassie Blue solution. Colonies containing more than 50 cells were counted, and the percentage of PGCs per colony was determined by dividing the number of PGCs by the total cell count in each colony. In the unirradiated controls, colonies were fixed and PGCs were calculated per colony at different stages of fBMSC clonal development. The same procedure was applied to measure the clonogenicity of SPSB-promoted MCF-10AS cells and breast cancer cells.

### fBMSC Proliferation Enhanced by PGCs

Due to different sensitivities to trypin between PGCs and proliferating diploid cells, the diploid cells in the colonies formed by juvenile phase fBMSCs were removed by a low concentration of trypsin (0.05% Trypsin-EDTA) applied at 37°C for 5 minutes. Subsequently, the remaining cells enriched with PGCs were collected through additional trypsinization using 0.25% Trypsin-EDTA. The collected PGCs were co-cultured with the same number of pre-stained juvenile phase fBMSCs (1×10^4^ CFSE-labeled cells per well) in a 12-well plate. The same number of unstained fBMSCs cocultured with pre-stained fBMSCs was included as the control. The coculture was maintained for 72 hours before quantification by flow cytometry for the pre-stained cells.

### mfPGCs

Mouse bone marrow-derived mesenchymal stem cells (mfBMSCs) were isolated from young (6-8 weeks) and old (24-27 weeks) female Balb/c mice and the mfBMSCs were collected from the femur and tibia aspirates using a standard method. The isolated mfBMSCs were washed twice with PBS and resuspended in BMSC maintenance medium and incubated for 72 hours and the adherent cells were washed twice with PBS. The medium of mfBMSCs was replaced every 3 days for a total of 10 days and then trypsinized and passaged up to 7 passages during which mPGCs were monitored.

### Biomarker of PGCs Spikings and SPSBs

IHC and confocal microscope were applied to detect an array of biomarkers including lysosomal, mitophagy, mitochondrial, EVs, or negative controls expressed on spikings and SPSBs in the dying-senescent PGCs of the phase III advanced-senescent fBMSCs. The phase III fBMSCs containing dying-senescent PGCs were living stained with mitoTrackerRed.

### SPSB Isolation and Purification

SPSBs were isolated from fBMSCs in advanced-senescent phase III maintained in T75 flask with a density of approximately 60-80% which contained about 30% PGCs by karyotyping and showed remarkable spiking beading (The cells were gently rinsed once with PBS to avoid disturbing the attached spikes. Subsequently, the cells were pretreated with 5 ml 0.05% trypsin at 37°C for 5 minutes to remove some loosely attached dead cells and debris. Then the pre-trypsinized cells were further digested with 5 ml 0.125% trypsin at 37°C for 5 minutes and stopped by adding an equal volume of exosome-free medium which selectively digests the attached SPSBs without inclusion of still attached cells. The collected SPSBs were suspended in 10 ml PBS and subjected to pre-centrifugation twice at 400×g and 450×g for 10 minutes each and the resulting pellets were then further centrifuged at 7000×g for 30 minutes. The purified SPSBs were resuspended in PBS and protein concentration was measured by BCA method, and aliquoted 100 µl into each Eppendorf tube and stored at −80°C.

### Ultrastructural analysis of SPSBs

Negative stain TEM grids were prepared by pre-fixing SPSBs and EXOs with 2% paraformaldehyde or 1% glutaraldehyde for 5 minutes, then placed in a droplet on parafilm and a copper formvar grid (Electron Microscopy Sciences, FCF200-CU-50) was floated on top for at least 30 minutes. Grids of SPSBS pre-fixed in paraformaldehyde were then transferred to a droplet of PBS, 2% glutaraldehyde, 8 consecutive water droplets, a droplet of 2% uranyl oxalate (pH 7) for 5 minutes, and a droplet of water. Grids of samples pre-fixed in glutaraldehyde were transferred to a droplet of water, then 2% uranyl acetate for 5 minutes. Grids from both methods were dried by dragging the edge across filter paper and leaving in air for at least 5 minutes to dry completely. Grids were imaged using the FEI L120C TEM. Cryo-EM grids were prepared using the FEI Vitrobot MkIII. Grids (Ted Pella, cat. 658-300-CU-100) followed by glow discharging, then loaded into the plunger apparatus. 4 μl of SPSBs was added directly to the grid and incubated for 20 seconds and the Grids were then autoblotted for 2 seconds before being plunged into liquid ethane for vitrification. Grids were stored in liquid nitrogen until imaging using the ThermoFisher Glacios.

### SPSBs mtDNA PCR

mtDNA enclosed within SPSBs was detected using mtDNA-PCR. To achieve this, 100 μl of purified SPSBs were boiled for 5 minutes, followed by centrifugation at 7000 rpm for 30 minutes. The resulting supernatant, containing mtDNA, was collected and utilized for PCR analysis. As a control, mtDNA samples were treated with DNase. For each reaction, 1 μl of mtDNA sample was employed along with specific pairs of mtDNA primers detailed below. The reactions were conducted using the Mastercycler PCR System (Eppendorf). Subsequently, the PCR products were identified through electrophoresis and sequencing to identify the mtDNA sequences.

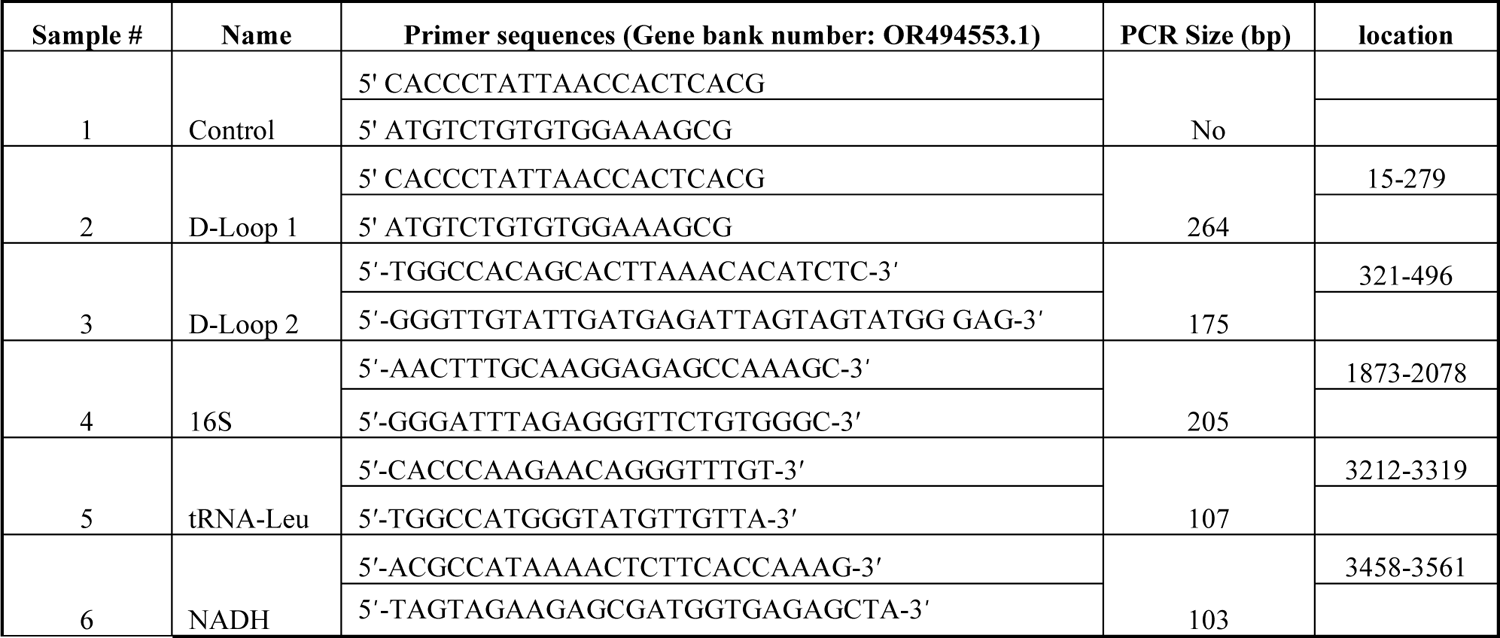

### THP-1 Cell Differentiation and Polarization

THP1 cells were treated with 2 μg/ml SPSBs for 24 hours, followed by induction of differentiation using 100 nM phorbol 12-myristate 13-acetate (PMA) for 48 hours. Subsequently, the cells were recovered in fresh culture medium for 72 hrs. Cells without SPSBs pretreatment were served as the control. For macrophage polarization, resting macrophages (M0) were primed with fresh medium supplemented with 20 ng/ml IFNγ and 100 ng/ml LPS for 48 hours to induce differentiation into the M1 phenotype. Similarly, to induce differentiation into the M2 phenotype, cells were primed with 20 ng/ml IL-4 and 20 ng/ml IL-13 for 48 hrs. The polarization was determined using flow cytometry. M1 phenotype cells were identified using CD80-FITC staining, while M2 phenotype cells were identified using CD206-APC-Cy7 staining.

### qPCR of SPSB-induced Macrophage Polarization

Total RNA was isolated utilizing the Trizol method followed by cDNA synthesis using the SuperScriptIII First-Strand kit. The primer pairs for detecting SPSB-mediated macrophage polarization are shown in the following chart by refereeing the reported results(64). Amplification reactions were performed using the Fast SyBR Green Master Mix with samples run on the CFX96 Touch Deep Well Real-Time PCR Detection System. The GAPDH gene was used as a housekeeping gene, and the results were analyzed using2^-ΔΔCT^ method.

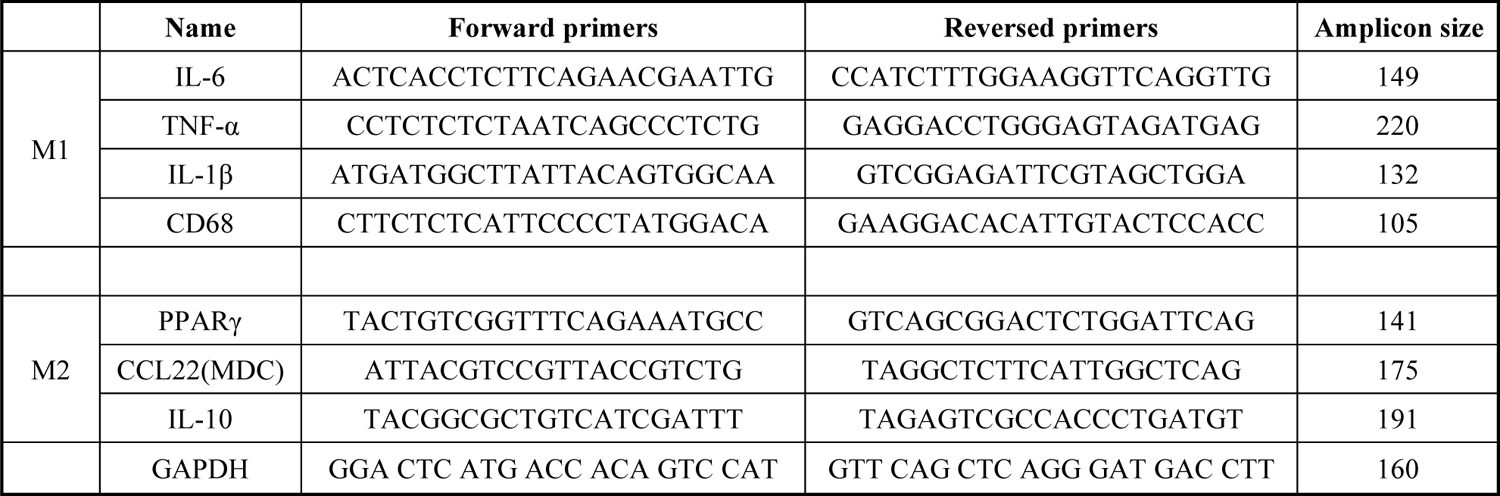

### SPSB-inhibited Macrophage Phagocytosis on Breast Cancer Cells

Phagocytic activity was conducted using differentiated macrophages derived from THP-1 cells with and without SPSBs endocytosis. In vitro phagocytosis assays were performed by pre-incubating THP-1 cells with SPSBs at a concentration of 2 μg/ml for 24 hours with untreated cells as the positive controls. The differentiated macrophages were generated by treatment with 40 nM Phorbol 12-myristate 13-acetate (PMA) for 48 hrs, then the polarized macrophages were cocultured at 37°C with GFP-labeled MCF-7 breast cancer cells at a 1:1 ratio for 2-4 hours. The cells were then collected by scraping, washed with 0.5% BSA-PBS, and incubated with fluorescence APC-Cyanine-7-labeled anti-CD11b antibody (BioLegend, 101225) in the dark for 30 minutes. After washing twice with 0.5% BSA-PBS, the phagocytic activity was analyzed using flow cytometry with the Becton Dickinson Canto II instrument (BD, NJ, USA) and data were analyzed with FlowJo VX.

### Nanoparticle Tracking Analysis

The size distribution of SPSBs and the control fBMSC-EVs that were generated from the culture medium of juvenile phase fBMSC and were characterized by using NTA following our precour publication (65). Both of the SPSBs and fBMSC-EVs were diluted in triple-filtered (0.22 μm) MilliQ-water to reach the working vesicle concentration. The diluted SPSBs and fBMSC-EVs were then analyzed using the NanoSight LM10 (Malvern Panalytical) equipped with a 404-nm laser and sCMOS camera. Three 90-s videos were recorded for each sample at camera level 12 using NTA v.3.0 software. Data were consistently analyzed with a detection threshold of 3 and screen gain of 10.

### Preparation of SPSBs Cargo Peptides

SPSBs peptides were prepared using the EasyPep Mini MS Prep Kit. The SPSBs pellets were dissolved in 100 µl of lysis buffer and protein concentration was measured by using the Pierce™ BCA Protein Assay Kit. Take 100 µg protein sample into a new microcentrifuge tube adjusted the final volume to 100 µl with a lysis solution. The sample was gently mixed with 50 µl of Reduction Solution and 50 µl of Alkylation Solution and incubated at 95°C for 10 minutes. After cooling to room temperature, the proteins were digested by adding 50 µl reconstituted enzyme solution and further incubated with shaking at 37°C for 3 hours before adding 50 µl stop solution. A volume ∼300 µl digested protein samples was then transferred into the dry peptide clean-up column and washed with 300 µl of wash solution A, followed by centrifuge at 15,000×g for 2 minutes, and then washed twice with wash solution B. The peptide clean-up column was then transferred into a new tube and centrifuged 1,500 ×g, 2 minutes to collect the cleaned peptides that were dried by a vacuum centrifuge and resuspended in 100 µl of 0.1% formic acid in water for LC-MS analysis.

### BMSCs-EVs Cargo Peptides

The juvenile fBMSCs were cultured with EXO-free medium for at least two passages, and the conditioned medium was collected and sequentially centrifuged at 300 xg for 10 min, 2,000×g for 20 minutes, and passed through a 0.22 mm filter. Then, the medium was concentrated using Amicon Ultra Centrifugal Filter Units with a 100-kDa MW cutoff (Sigma), transferred to thick wall polypropylene tubes (Beckman Coulter), and centrifuged at 8,836×g using the SW28 rotor and L7 Ultracentrifuge (Beckman Coulter). The supernatant was transferred to fresh tubes, and centrifuged at 112,700×g for 90 minutes which was repeated once, the final EVs pellet was resuspended in PBS, identified with EVs biomarkers, and stored in aliquots at −80^◦^C.

### LC-MS of Cargo peptides of SPSBs versus BMSC-EVs

Liquid chromatography was performed at 40°C on ultra-high-pressure nano-flow Easy nLC (Bruker Baltonics) with a constant flow of 400 nl/minutes on a PepSep 150µm×25cm C18 column (PepSep, Denmark) with 1.5 μm particle size (100 Å pores) and ZDV spray emmiter (Bruker Daltronics). Mobile phases A and B were water with 0.1% formic acid (v/v) and 80/20/0.1% ACN/water/formic acid (v/v/vol), respectively. Mass spectrometric analysis was performed on a hybrid trapped ion mobility spectrometry-quadrupole time of flight mass spectrometer (*timsTOF Pro*, (Bruker Daltonics, Bremen, Germany) with a modified nano-electrospray ion source (CaptiveSpray, Bruker Daltonics) and operated in PASEF mode. Data-independent analysis (DIA) was performed on a nanoElute UHPLC coupled to a timsTOF Pro. The acquisition scheme used for DIA consisted of four 25 m/z precursor windows per 100ms TIMS scan and 16 TIMS scans, creating 64 total windows and layered the doubly and triply charged peptides on the m/z and ion mobility plane. Precursor windows began at 400 m/z and continued to 1200 m/z and the collision energy was ramped linearly as a function of the mobility from 63 eV at 1/K0=1.5 Vs cm^−2^ to 17 eV at 1/K0=0.55 Vs cm^−2^.

### Proteomics Analysis

The MS raw files were processed with Spectronaut, v.16.3, and all searches applied the protein sequence database of reviewed Homo Sapiens proteins UP00000 with decoys and 115 common contaminant sequences. Decoy sequences were generated and appended to the original database with a maximum of two missing cleavages allowed, the minimum peptide sequence length as 7 amino acids, and the peptide mass limited to a maximum of 4600 Da. Carbamidomethylation of cysteine residues was set as a fixed modification, and methionine oxidation and acetylation of protein N termini as variable modifications. The initial maximum mass tolerances were 70 ppm for precursor ions and 35 ppm for fragment ions. A reversed sequence library was generated/used to control the false discovery rate (FDR) at less than 1% for peptide spectrum matches and protein group identifications. Decoy database hits, proteins identified as potential contaminants, and proteins identified exclusively by one site modification were excluded from further analysis. The list of quantified proteins was uploaded to SimpliFi (ProtiFi, LLC), an online platform for statistical analysis and visualization of omics data.

### Bioinformatics Analysis

The biological process-, cellular component- and molecular function-related analysis of the genes corresponding to the proteins enriched > 4 folds in SPSBs compared to MSC-EXO were performed using ShinyGO 0.77 software (http://bioinformatics.sdstate.edu/go/). For the bioinformatic analysis of human tissues and cancer patient data, normalized read counts for genes expression in patient tissues and patient clinical information in TCGA datasets were downloaded from the UCSC xena [http://xena.ucsc.edu/]. Samples were ranked using each normalized gene expression. Overall survival analyses were performed for the high and low expression ranked values for *AIFM1, NDUFAB1, NDUFS1*, *PPA2*, *CDH13*, *ERCC6L*, *ACSF2*, *ARID5A*, *CHI3L1*, *CXCL3*, *CXCL5*, *CXCL16*, *CYTIP*, *FHIT* and *IZUMO4* and plotted for Kaplan-Meier curves using GraphPad Prism (GraphPad).

### Breast Cancer Cohort Analysis

The Kaplan-Meier Plotter online software (http://kmplot.com/) was used to analyze the relationship between the OXPHOS gene expression and the prognosis of breast cancer patients. Two cohorts with high and low expression of each of OXPHOS genes were compared by Kaplan-Meier survival analysis, and hASrd ratios and logrank *P*-values with 95% confidence intervals were calculated. Public database platform TIMER (https://cistrome.shinyapps.io/timer/) were used to compare the expression level of each of OXPHOS genes between breast cancer and surrounded normal breast tissues using the TCGA database. The distribution of gene expression levels was displayed using boxplots and the statistical significance of differential expression was assessed using the Wilcoxon test. Public database platform Sangerbox (http://sangerbox.com/home.html) were used to compare the difference between breast cancer with normal tissues and their related prognosis of breast cancer in TCGA database. Kaplan-Meier survival analysis was applied to compare two cohorts of breast cancer with high and low expression of OXPHOS genes, calculating the hASrd ratios and log rank P values with 95% confidence intervals.

### Transcriptomics of SPSB-promoted MCF-10AS

MCF-10AS and the control counterpart MCF-10A cells with similar passaging were pelleted by centrifugation and stored at −80°C. Total cellular RNA was isolated using the TRIzol reagent (Invitrogen, 15596-018) and also a modified protocol that incorporates an additional extraction with phenol/chloroform/isoamyl alcohol (25:24:1, pH 4.3) was included. RNA quantity and quality were assessed on a NanoDrop spectrophotometer (Thermo Scientific) and the Agilent 2100 Bioanalyzer (Agilent Technologies), respectively. Then total RNA was converted to cDNA and used as the input for a next-generation sequencing library preparation. RNA-seq was performed for transcriptome profiling, differential gene expression, and functional profiling. Reads were separated by human using Xenome, and they were next aligned and quantified using TopHat2, Cufflinks, hg38 and UCSC transcript and gene definitions.

### SPSB-promoted Breast Epithelial Cell Protumorigenesis

Human breast epithelial MCF-10A cell line was subjected to the test of SPSBs-promoted breast epithelial cell protumorigenesis. MSC-10A cells in completed medium were constantly exposed to SPSBs with a concentration of 4.4×10^6^/cell in 60 mm cell culture dishes and SPSBs was refreshed at each passage and the coculture was maintained for at least 6 passages and the resulted MCF-10AS cells were applied for the studies of SPSBs-mediated transformation. SPSB-promoted MCF-10A cells were served as the control counterpart. For protumorigenic analysis, MCF-10AS and the control MCF-10A cells were investigated with the acquired aggressiveness including clonogenicity, wounding healing, 3-D culture, matrigel invasion and in vivo mammary plug acrinol structural analysis.

### Wound Healing Assay

A wound healing assay was used for revealing the cell migration capacity of MCF-10AS and the counterpart control MCF-10A cells. 1 × 10^6^ cells of each cell type were seeded into 6-well plates. When reached 100% confluence, the medium was replaced with 1% FBS starvation medium, and cells were further cultured for 24 hours before the gap was created by scraping the dish diagonally with a sterile P200 tip. The cell migration capacity was monitored at 0-, 12-, 24-, and 48-hours post scraping.

Images were obtained by phase contrast microscopy, and the gap-filling ability of cells was estimated by measuring the gap distance and quantitation using Image J (NIH Image).

### Transwell Invasion

Matrigel (BD Biosciences, 356231) was diluted with the coating buffer: 0.01 M Tris (pH 8.0), 0.7% NaCl at final concentration of 200-300 µg/ml (1:40-45 dilution from stock) and 100 μl of the diluted matrigel was added into upper chamber of 24-well transwell (Corning, 3422) and incubated at 37°C for 2 hours for gelling. The coating buffer was then removed from the permeable support membrane and the cells to be tested were resuspended in medium containing 1% FBS at a density of 2.5 ×10^4^ cells/300 μl before adding onto the upper chamber. The lower chamber was then filled with 800 μl of fresh cell culture medium containing 5 μg/ml fibronectin (Corning, SC-29011, x). Cells were incubated for 48 hrs and the cell penetrating capacity was measured with the membrane stained with Diff-Quick Stain kit (IMEB INC, K7128) and analyzed and quantitated by ImageJ.

### 3-D Mammosphere Analysis

The three-dimensional culture of MCF-10A cells was conducted following the standard method. Briefly, 40 μl Growth Factor Reduced Matrigel (GFRM) was added to each well of an 8-well LabTek Chambered cover glass and incubated at 37°C for 30 minutes, followed by adding 5,000 cells in 200 μl GFRM supplemented with 5 ng/ml EGF, 2% (v/v) FCS, 4% (v/v) Matrigel. The culture medium was changed every 3 days until the morphology was assessed after 10 days. The 3-D acinus was then fixed, and the IF images were captured with confocal microscopy ZEN3.6 (ZENpro) using ZEISS ZEN3.6 software. The laminin V antibody staining was included to indicate basement membrane deposition for SPSB-promoted MCF-10AS and control cells.

### In vivo Mammary Acinus Morphogenesis

The MCF-10AS cells and the control MCF-10A cells were transduced with lentivirus vectors carrying GFP-Luciferase. Subsequently, 1 × 10^6^ cells suspended in 100 μl of matrigel were implanted into the mammary fat pads of 6-week-old female NRG mice, and tumorigenesis was monitored for three weeks. The mammary fat pads were surgically removed, fixed, and embedded in paraffin. Paraffin sections measuring 7 μm in thickness were prepared following the standard immunohistochemistry protocol. The primary antibody mixture, consisting of anti-GFP and anti-laminin V, was dissolved in 1% (v/v) BSA in PBS and incubated overnight at 4 °C. After washing with PBS, a secondary antibody mixture containing Alexa Fluor 488-conjugated mouse secondary antibody and Alexa 594-conjugated rabbit secondary antibody was incubated for 1 hourat room temperature. The slides were then mounted using DAPI mounting medium. Imaging was performed using ZEN3.6 (ZENpro) microscopy and processed using ZEISS ZEN3.6 software.

### Identification of PGCs in Mouse Spontaneous Breast Tumor

Generation of mouse spontaneous breast tumor was followed by the established protocol(66). C57BL/6 mice (6–8 week old female 20-25 g weight) were purchased from Jackson Laboratory (No: 002019) and were gavaged weekly for six consecutive weeks with 1 mg/100μl DMBA dissolved in olive oil. Starting from the third week of DMBA administration, 2 mg/50μl MPA was subcutaneously injected every two weeks for a total of 3 treatments. At the end of the administration, mice were monitored for body weight change and tumor formation twice a week for calculating the tumor volume = tumor length × tumor width^2^/2. When the tumor volume of experimental mice reaches to 1400 mm^3^. Mice were euthanized and tumors were collected for pathological identification and PGC analysis.

### SPSB Biomarker in PGCs in SPSB-promoted MCF-10AS and BC Tissues

The PGC formed in the SPSB-promoted MCF-10AS, BC cells and tissues of BC and mouse spontaneous breast tumors were identified following the standard of pathological definition of PGCs that contain a large or multiple nuclei and are at least three times larger in size as compared with that of parental cancer cells (37, 49, 67). Pathologically diagnosed BC subtypes in Hematoxylin-Eosin (H&E)-stained slides from UC Davis Comprehensive Cancer Center Biorepository were evaluated using standard light microscopy. PGCs in SPSB-promoted MCF-10AS cells and BC cells were identified by karyotyping and calculated in the growing edge of wound healing of MCF-10AS cells and colonial expansion of BC cells to reveal the percentage of PGCs during cell proliferation and clonal expansion.

## Supporting information

SuppFigs

## Data Availability

The data that support the findings of this study are within the Article, Supplementary, including Figures and videos, or available from the corresponding author upon reasonable request.

## Authors’ Contributions

J.J.L., B.X., F.A.F., L.J., and M.F. conceived the project. B.X., F.A.F. and M.F. maintained and characterized human BMSCs. B.X, C.X.W., M.F., S.X., Y.D., J.H., Y-Y. Z., J.I.B., D.W., A.J., A.W. and J.J.L. conducted experiments and data analysis on BMSCs and SPSB-promoted breast epithelia cells. M.F., Y.Z., C.X.W., T.-y.L., Y.L. D.H., and A.W. conducted MSC IHC analysis. D.H., K.G., R.M. and R.C. conducted and analyzed TEM ultrastructure data. B.W., S.X. and M.F. conducted experiments and data analysis in macrophage and breast epithelial cells. Q.G., L.J. and J.J.L conducted and analyzed the living cell dynamics. B.X., Y.D., M.F., Y.S., M.D., A.M.M., C.X.W. and S.X. conducted mouse in vivo experiments. B.X., C.F., S.X., and M.F. prepared the sample and analyzed the proteomics and transcriptomic data. Y.Z. conducted the mouse breast tumor and breast cancer pathological analysis. B.W., M.F., L.J., A.M.M., F.A.F., and J.J.L. wrote the manuscript.

## Acknowledgements

This work was supported by National Cancer Institute Grant R01 CA213830 (JL), University of California Davis Cancer Center Cancer Immunology Pilot Grant Support (JL), and the University of California Davis NCI-designated Comprehensive Cancer Center supported by the CCSG Grant awarded by the National Cancer Institute (Dr. Primo Lara; NCI P30CA093373). We thank Dr. Xiao-Jing Wang at UC Davis Cancer Center for recording live-cell imaging; Dr. Gabriela Grigorean at UC Davis Proteomics Core Facility for assistance in SPSBs cargo protein preparation and LC-MS/MS data analysis; Dr. Regina Gandour-Edwards at UC Davis Pathology Biorepository for providing breast cancer pathological samples. We acknowledge the service of UC Davis Cancer Center Research-Supporting Cores for cellular and animal studies.

