## Supplementary material for "Post-death Vesicles of Senescent Bone Marrow Mesenchymal Stromal Polyploids Promote Macrophage Aging and Breast Cancer": SuppFigs

**Supplementary Figures: S**1-10
**Supplementary Videos: S**1-S2

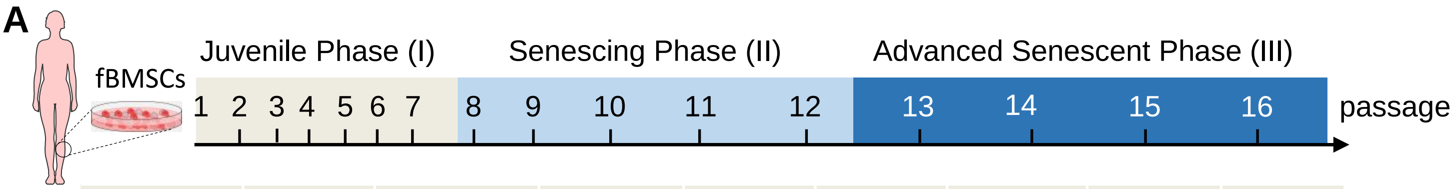


| Donor Code | 20 | 22 | 43 | 52 | 60 | 63 | 66 | 68 |
|---|---|---|---|---|---|---|---|---|
| Gender | F | F | F | F | F | F | F | F |
| Age | 44 | 45 | 32 | 49 | 52 | 52 | 35 | 19 |
| Health Condition | Healthy | Healthy | Healthy | Healthy | Healthy | Healthy | Healthy | Healthy |

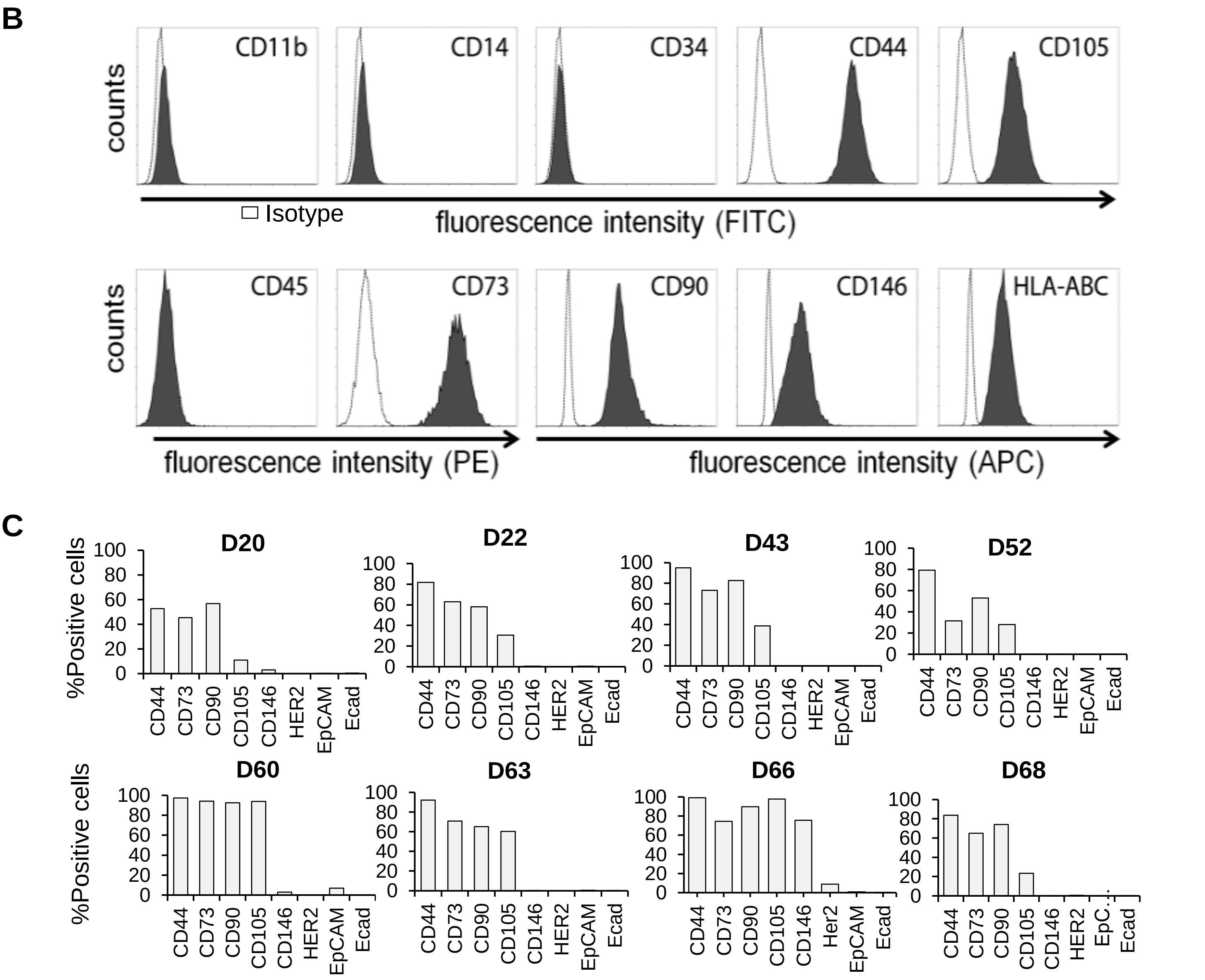


**Supplementary Fig. S1.** fBMSCs from healthy female bone marrow donors identified by MSC and non-MSC biomarkers**. A**, Schematics of senescence course induced by in vitro expansion of fBMSCs from a group of healthy female bone marrow donors (n = 8). **B**,**C**, Representative flow cytometry of positive human MSC biomarkers (CD44, CD73, CD90, CD105, CD146, HLA-ABC) compared to the negative control biomarkers (CD11b, CD14, CD34, CD45) and a group of non-MSC control receptors (HER2, EpCAM, and Eoad).

#### Supplementary Fig. S2

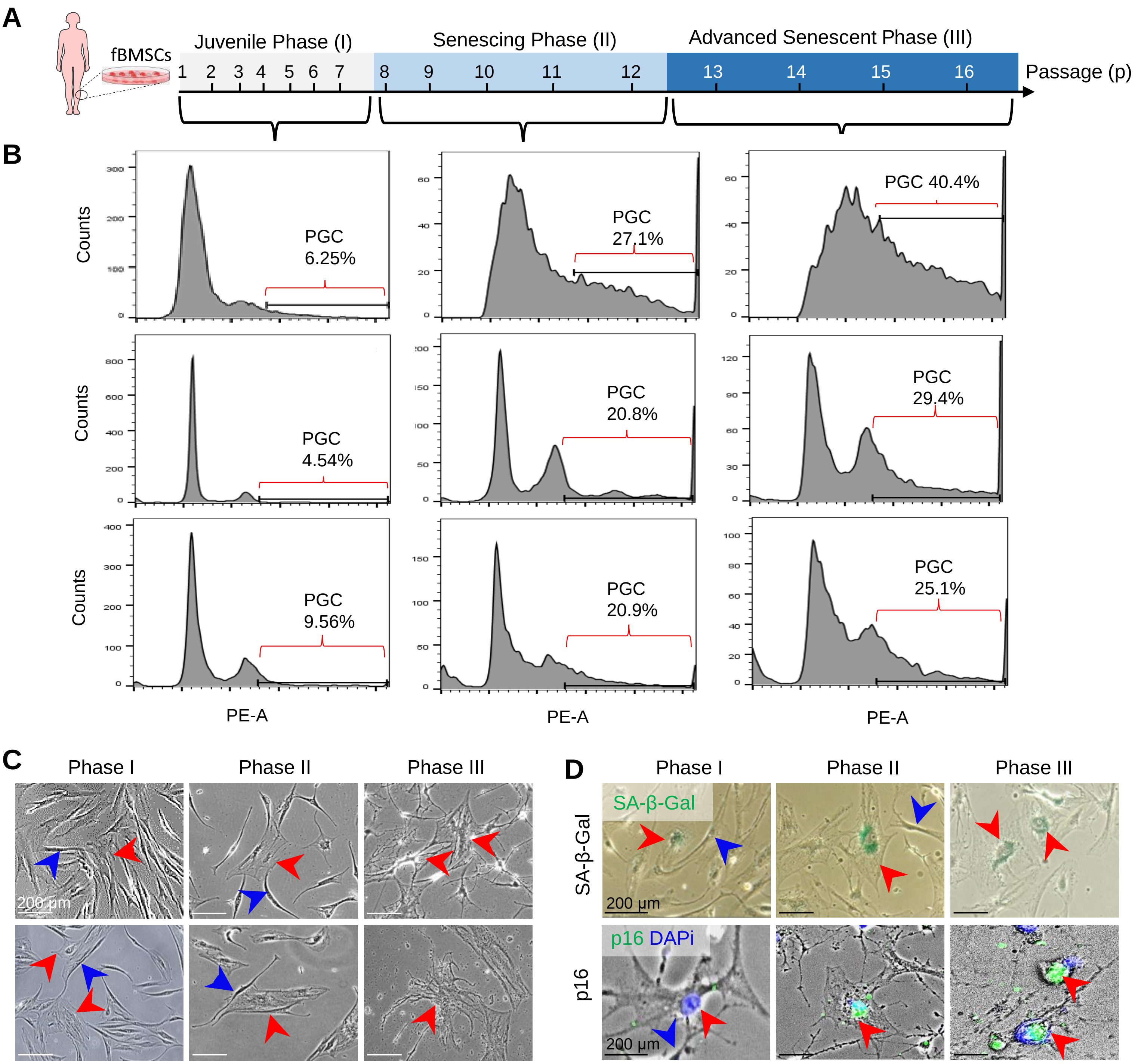


**Supplementary Fig. S2.** Karyotyping, morphologies, and senescence of PGC in juvenile (phase I), senescing (phase II), and advanced-senescence (phase III) fBMSCs. **A**, Schematic of karyotyping PGCs in the indicated phases of fBMSCs. **B**, Representative PGC populations in three sets of fBMSCs calculated by using the baseline of juvenile phase fBMSCs as > 4n genome from the same fBMSC donor. **C**, Representative PGC morphologies in the indicated senescent phases of fBMSCs from two BM donors (red arrow). The blue arrow indicates the fBMSCs attached to PGCs in the juvenile and early senescing phases. **D**, Representative senescent PGCs (red arrow) stained with SA-β-galactosidase and p16 (green) with nucleus stained with DAPi (blue). The blue arrow indicates the fBMSCs attached to PGCs in the juvenile and early senescing phases.

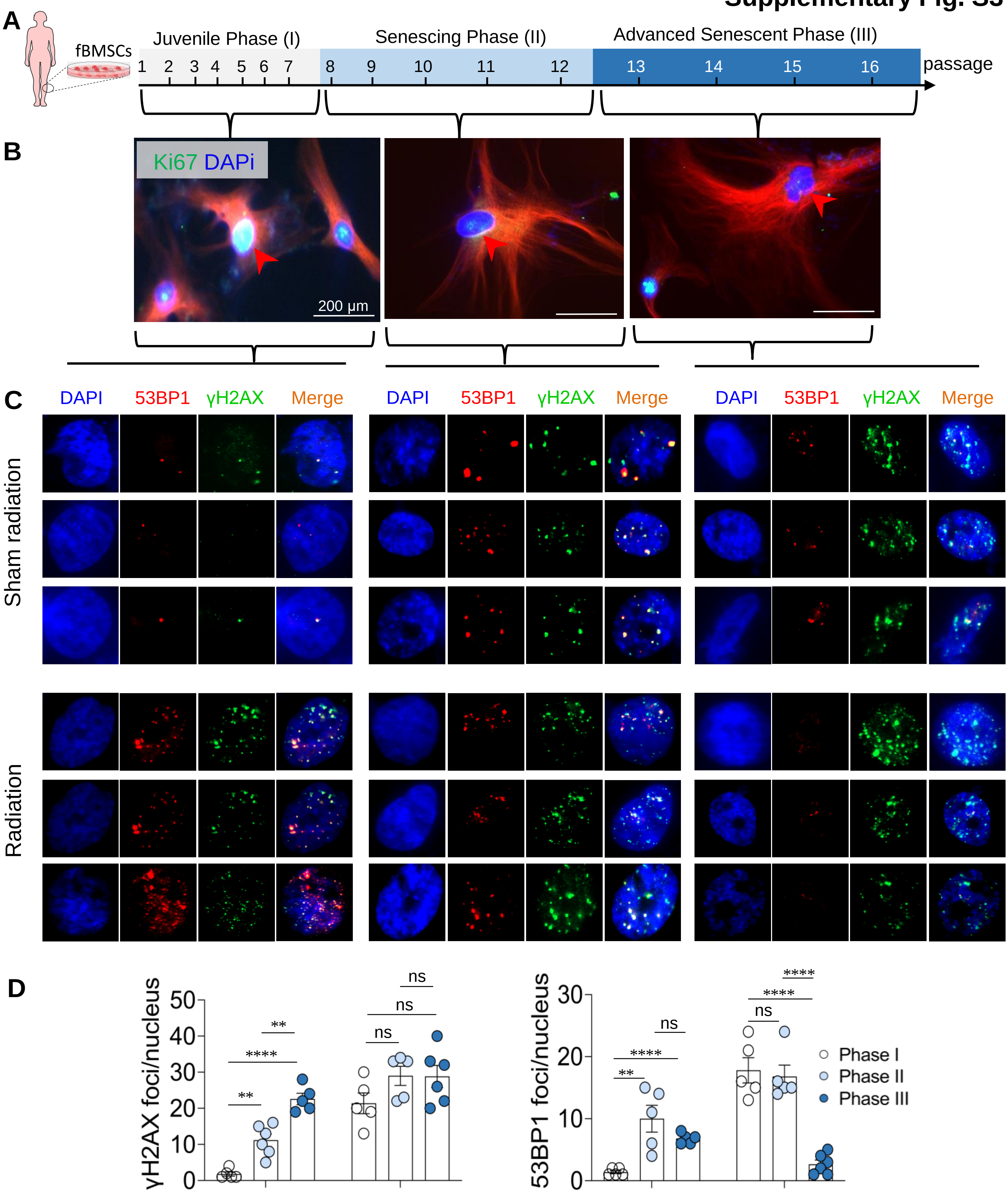


**Supplementary Fig. S3.** Proliferative and DNA repair capacity of PGCs**. A**, Schematic of proliferative and DNA repair activity measured in the indicated senescent phases of fBMSCs**. B**, Representative images of PGCs in the indicated fBMSC senescent phases stained by proliferative biomarker Ki67 (green; nucleus stained by DAPi). Images (**C**) and quantitation (**D**) of DNA repair activity of PGCs at indicated fBMSCs senescent phases measured by γH2AX and 53BP1 foci formation 72 hours after irradiation with a single dose of 2 Gy x-ray. Sham irradiation was used as a control. n = 6, ** $p< 0.01$. *** $p< 0.001$. **** $p< 0.0001$, ns, $p> 0.05$.

### Supplementary Fig. S4

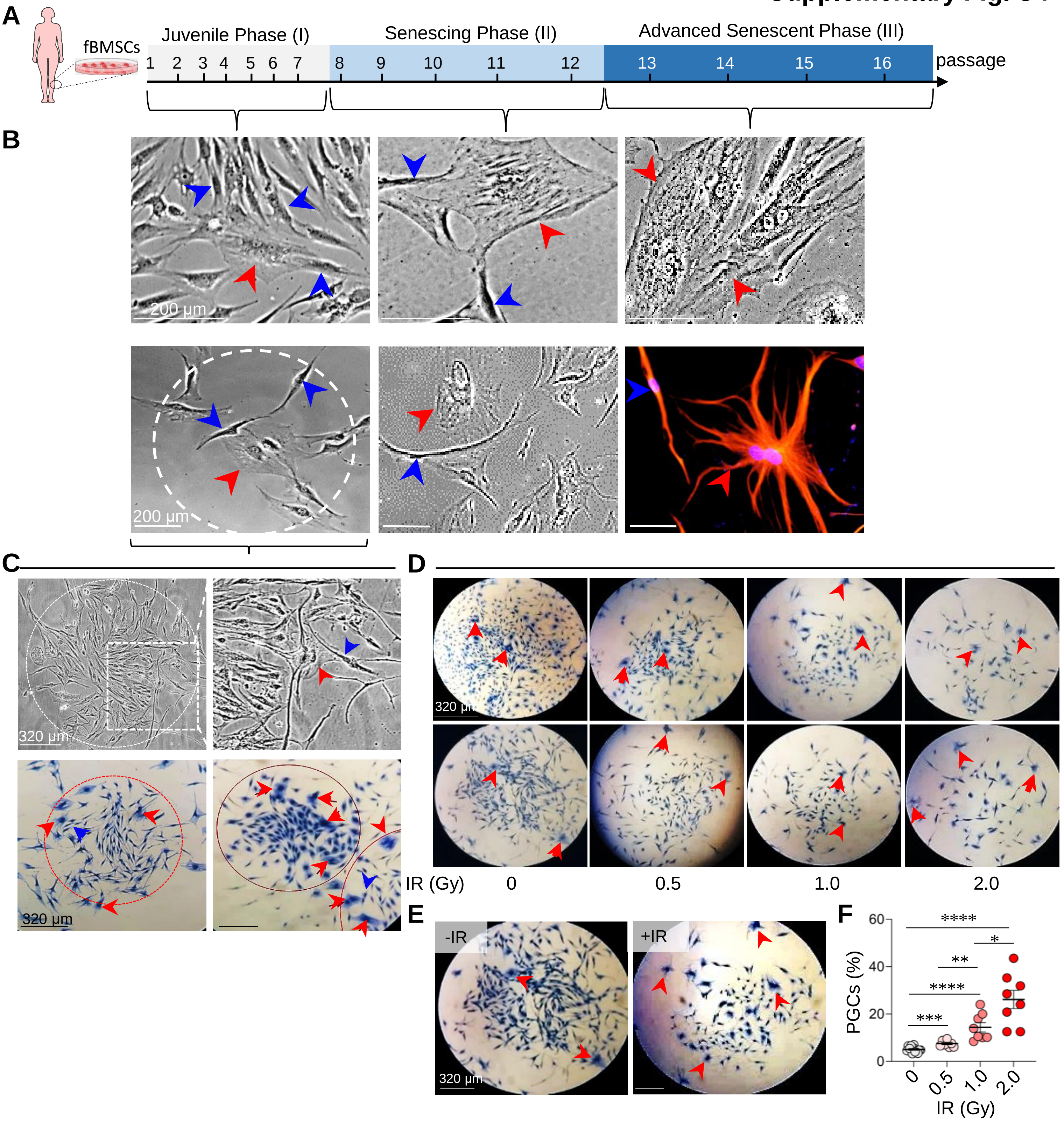


**Supplementary Fig. S4.** PGC-associated fBMSC clonal expansion and radiation-surviving colonies. **A,** Schematic time of monitoring PGCs in boosting neighboring cell clonal expansion**. B,** Representative living PGCs (red arrow) interacting with neighboring cells (blue arrow) in phase I and II fBMSCs compared to the absence of such activity with the senescent PGCs in phase III fBMSCs labeled with RFP. **C,** Living (up panel) or fixed (lower panel) colonies generated from phase I fBMSCs (red arrow, PGCs located in the growing edge of the colony; blue arrow, proliferative cells attached to PGC). **D**, Representative images of PGCs (red arrow) in fBMSC colonies surviving radiation of indicated doses. **e**, Representative fBMSCs colonies with or without radiation indicating an increased PGC fraction in the radiation-surviving colony. **f**, Percentage of PGCs per colony in radiation-surviving fBMSC colonies (n = 7; **p*< 0.05, ** *p*< 0.01, ****p*< 0.001, *****p*< 0.0001, ns, *p*> 0.05 , data are shown as mean ± SEM).

#### Supplementary Fig. S5

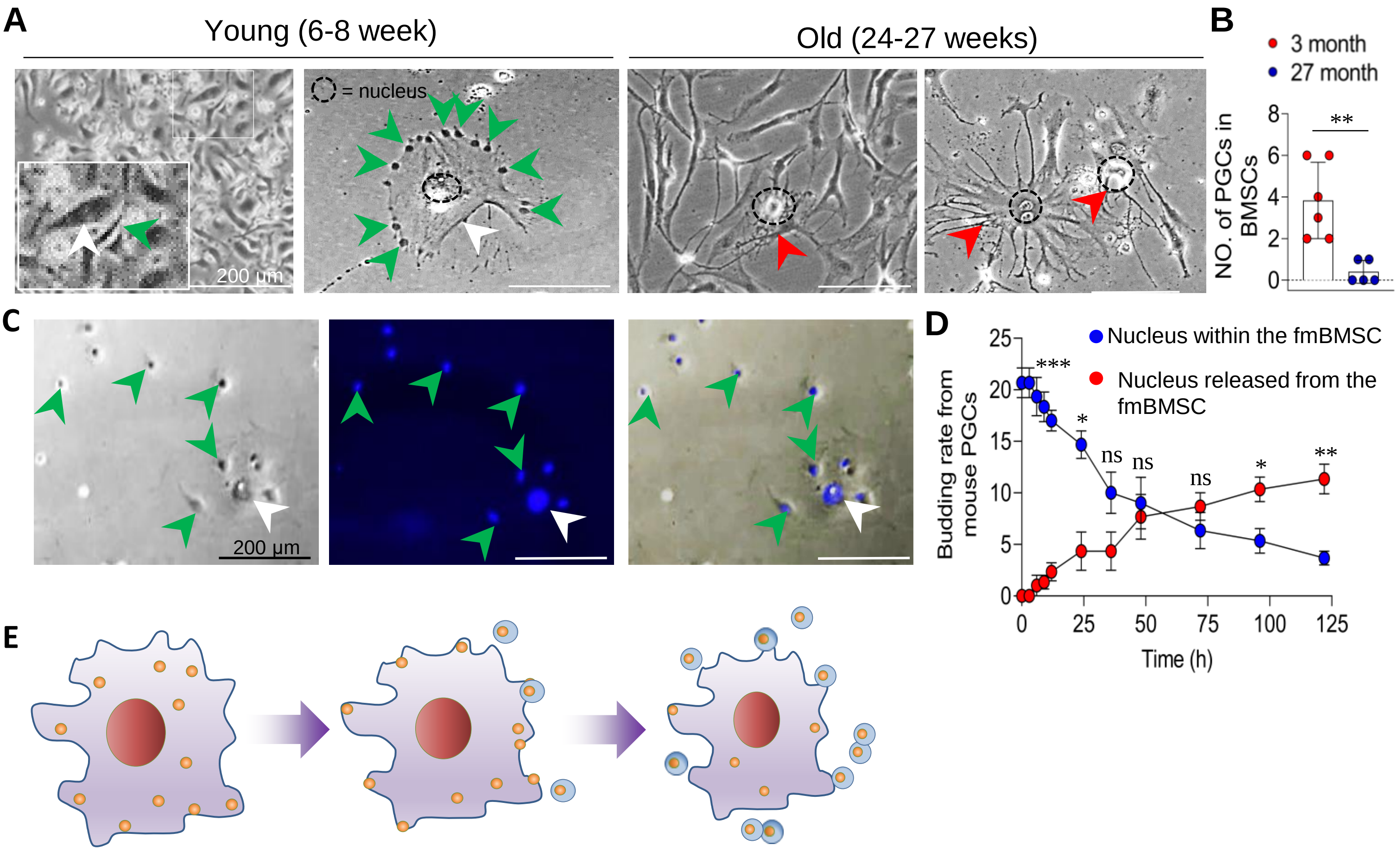


**Supplementary Fig. S5.** fmPGC boosts diploid cell proliferation. **A**, Living images of fmPGC attached with diploid cells and dandelion-seeding-like cell budding from an fmPGC detected in fmPGC from young (6-8 week) mice which were not detected in fmBMSCs from old (24-27 weeks) mice. White and red arrows indicate the juvenile and senescent fmPGCs respectively; green arrows indicate the diploid cell or the nucleus attached to the inner membrane of fmBMSCs. respectively. **B**, Quantification of dandelion-seeding-like budding cells in young and old fmBMSCs (young mice n = 6; old mice n = 5, ** *p* <0.01). **C**, Living image of budding cells (green arrows) from an fmPGC (white arrow) with nuclear DNA of both the fmBMSC and budding cells visualized by DAPi (green arrow). **D**, Time course of cell budding calculated by counting the nucleus within (blue) and released (red) from the fmBMSC. n = 5, **p*< 0.05, ***p*< 0.01, ****p*< 0.001, ns, *p*> 0.05. **e**, Schematic model of fmPGCs promoting cell proliferation via dandelion-seeding-like cell budding.

### Supplementary Fig. S6

A

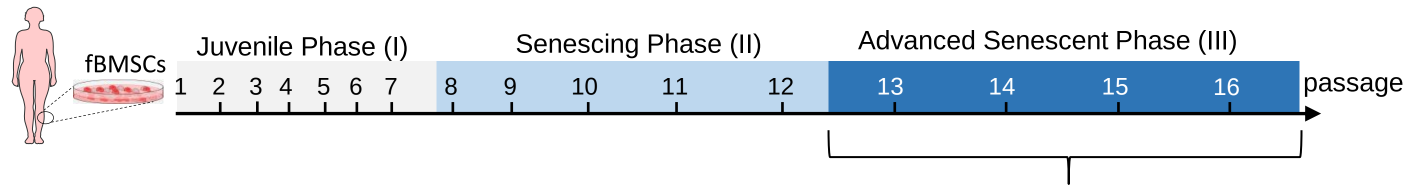


B

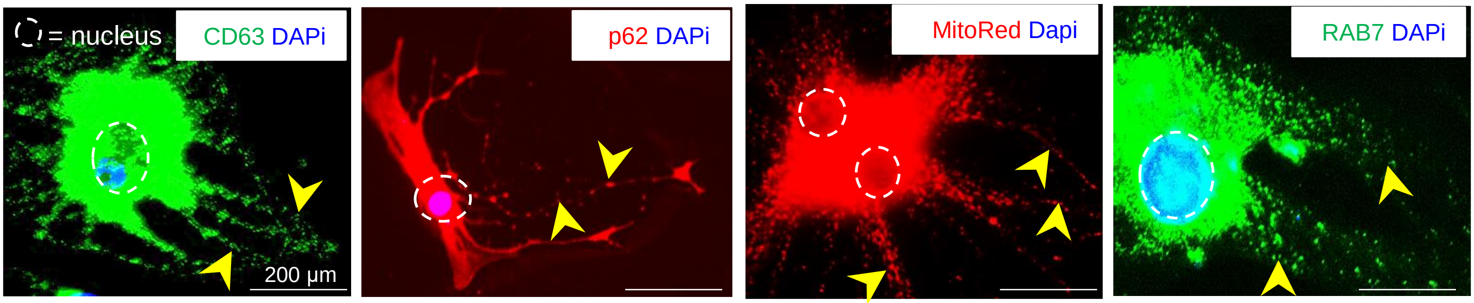


C

**Summary of SPSB Biomarkers**

| Markers | Vesicles/organelles | Relative expression |
|---|---|---|
| CD9 | EVs | (-) |
| CD81 | EVs | (-) |
| Tublin | cell skeletal system | (+) |
| RAB7 | late endosomes | (+++) |
| p62 | mitophagy | (+++) |
| Lamp1/CD63 | lysosome/endosome marker | (+++) |
| LysoTracker | lysosome | (+++) |
| MitoTracker | mitochondria | (+++) |

D

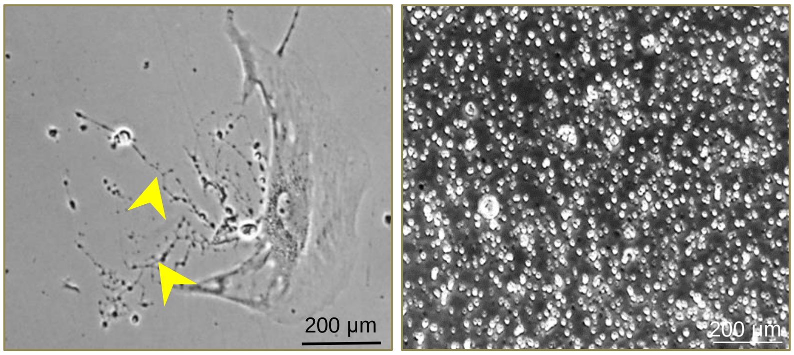


E

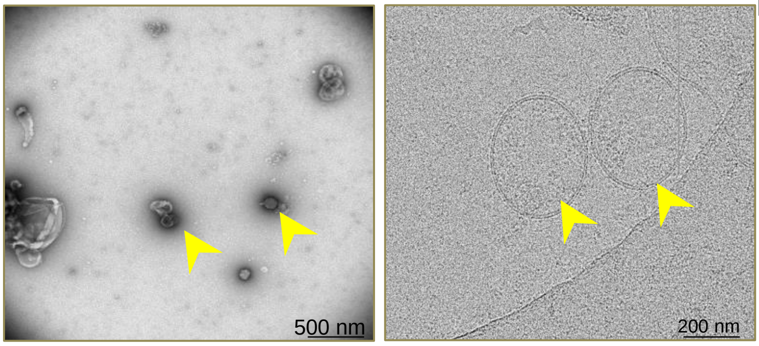


F

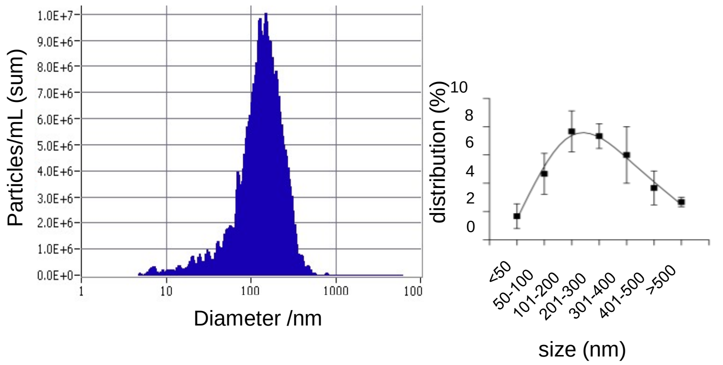


G

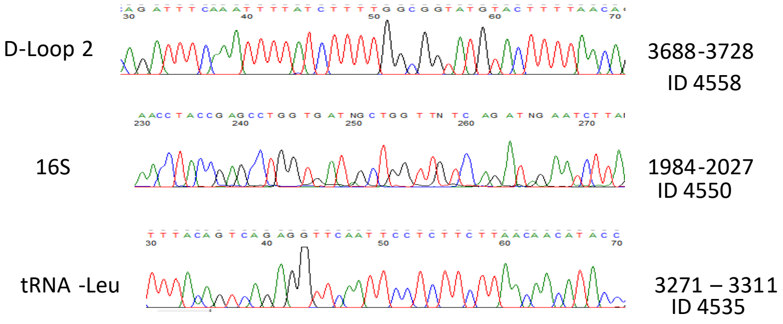


**Supplementary Fig. S6.** Characterization of SPSBs**. A.** Schematic period of phase III fBMSCs used for isolation of SPSBs from dying PGCs. **B**, Representative IF images of SPSBs (yellow arrows) separating from dying PGCs stained with lysosomal and mitophagosome biomarkers. **C**, Summary of tested biomarkers. **D,** Representative images of SPSBs before (left) and after (right) purification. **E**, Images of SPSBs by TEM with negative stain (left) and Cryo-EM (right). **F,** SPSBs volume distribution measured by NTA (left) and by TEM (right). **G,** Identification of mtDNA sequence in isolated SPSBs using three pairs of mtDNA primers (D-Loop, 16S, and tRNA-Leu).

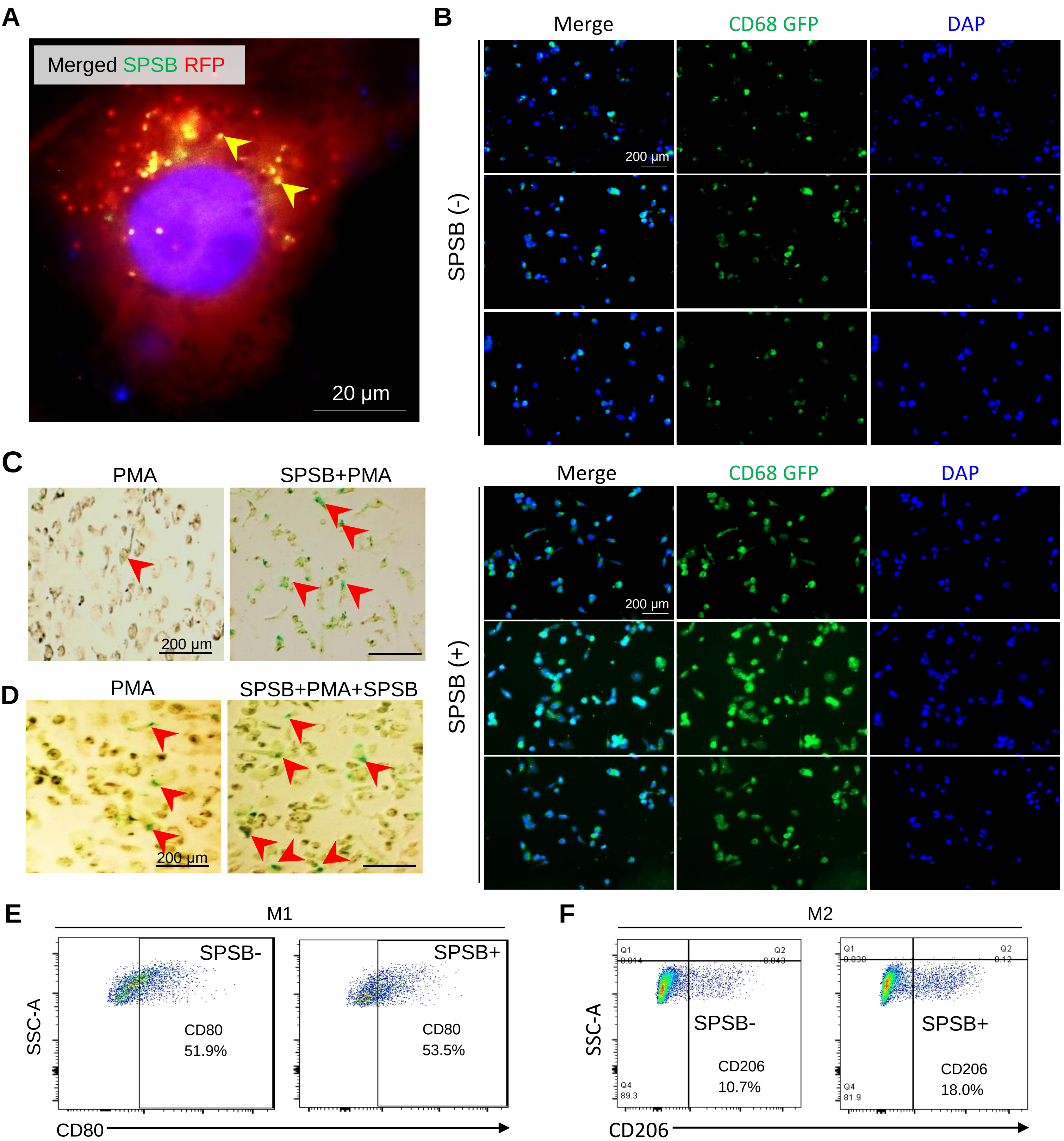


**Supplementary Fig. S7.** SPSBs boost macrophage maturation, senescence, and protumor M2 polarization. **A**, SPSBs (GFP, yellow arrows) endocytosed by an RFP-macrophage (nucleus = blue by DAPi). **B**, Representative IF images of CD68-expressing macrophages differentiated by PMA with or without pretreatment with SPSBs. Representative images of senescent macrophages stained by β-gal after differentiation by PMA or SPSB+PMA for 72 hours (**C**) or continued for an additional 72 hours (**D**) with PMA or SPSB+PMA (red arrows indicate the senescent macrophages). Flow cytometry analysis of M1 (**E**) and M2 (**F**) polarization of macrophages differentiated from PMA and SPSB+PMA treated THP-1 cells from d.

**Supplementary Fig. S8**

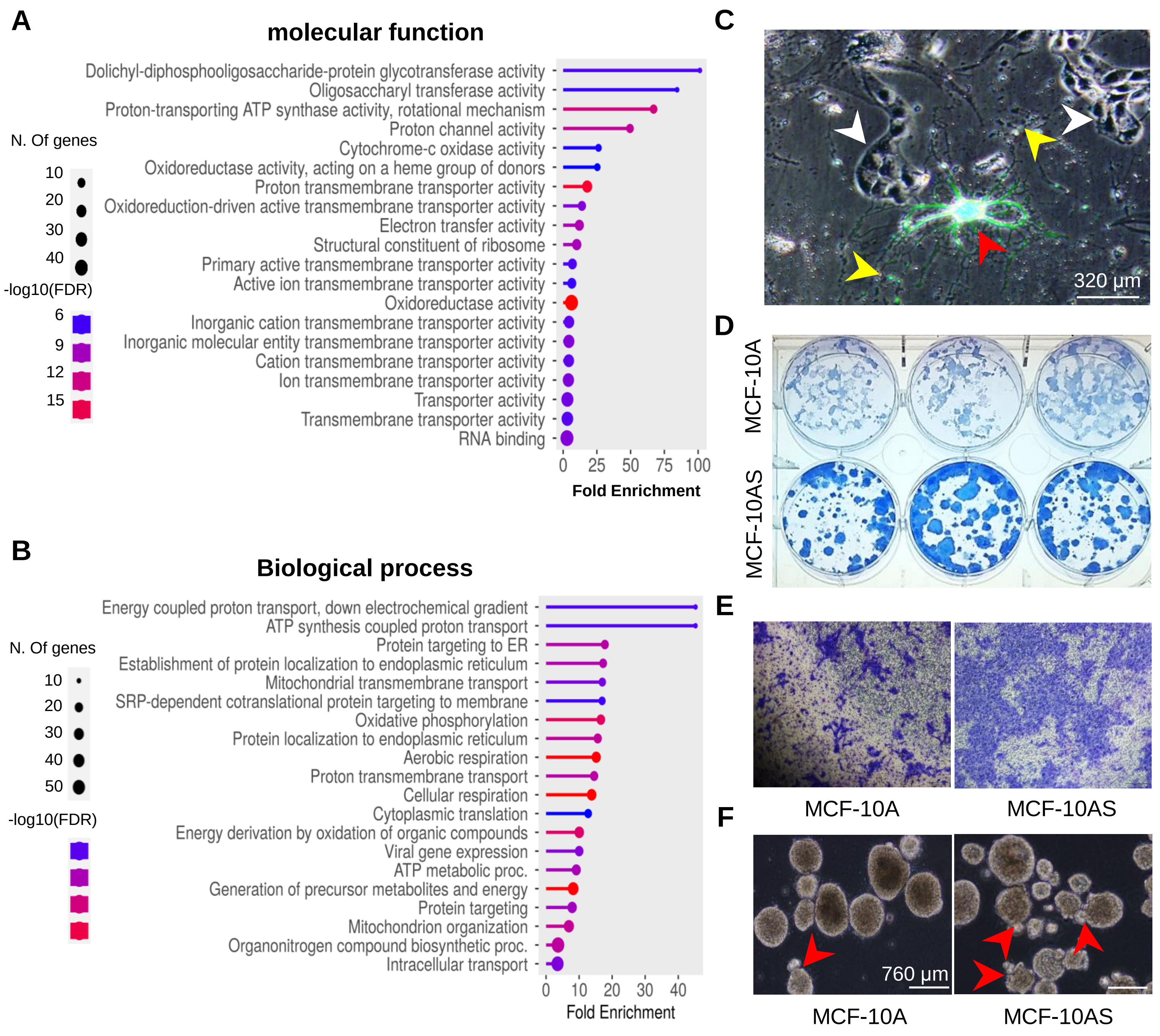


**Supplementary Fig. S8.** Proteomics of SPSB cargo versus BMSC-EVs and protumorigenic phenotype of SPSB-promoted MCF-10AS cells. Functional (**A**) and related biological network (**B**) of proteins enriched in SPSB cargo compared to hBMSCs-EVs by proteomic analysis. **C**, MCF-10A clonal formation (white arrows) attached to a senescent GFP-PGC (red arrow) that releases SPSBs (yellow arrow) in the coculture of MCF-10A cells with advanced-senescent GFP-fBMSCs. Representative images of clonogenic MCF-10A and SPSB-incorporated MCF-10AS cells that demonstrate enhanced clonogenicity (**D**), Matrigel penetration (**E**), and phase-contrast morphology of 3-D acini formation (**f**, Red arrows indicate protumorigenic acini with protrusions in MCF-10AS cells).

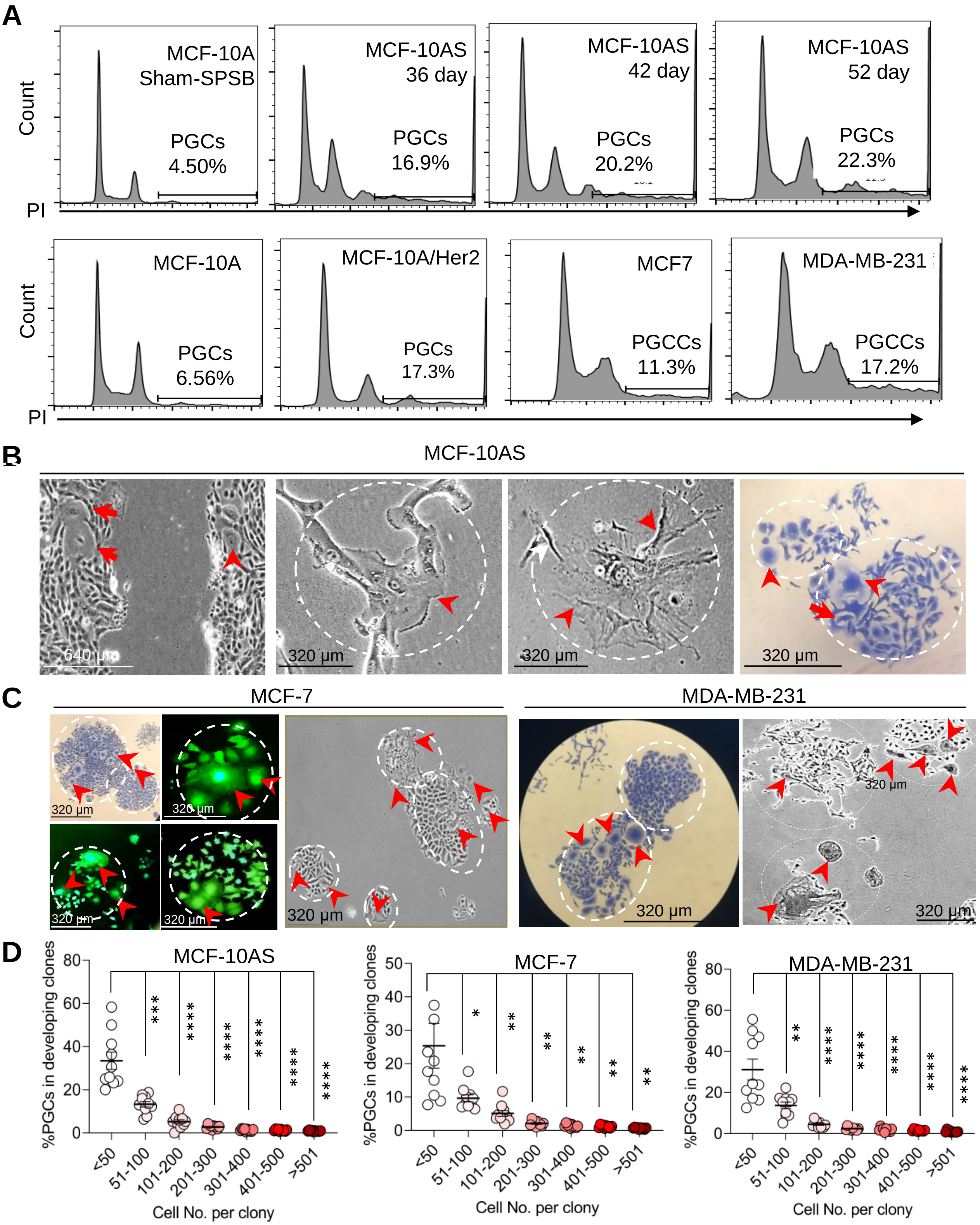


**Supplementary Fig. S9.** PGCs and PGCCs in MCF-10AS and breast cancer cells. **A,** Karyotyping of MCF-10AS cells generated from coculture of SPSBs for indicated periods compared to the basal PGCs in MCF-10A cells under sham-SPSB culture condition which was compared to the PGCs in the parental MCF-10A cells with or without transformation by overexpressing Her2 gene, and the PGCCs in BC cells MCF-7 and MDA-MB-231 cells. **B**, Representative image of PGCs in the growing edge of gap filling and in starting and established colonies of MCF-10AS cells**.** **C**, Images of PGCCs in different stages of colonies generated by MCF7 and MDA-MB-231 cells. **D**, Percentage of PGCs/colony was similarly decreased with clonal expansion of MCF-10AS, MCF-7, and MDA-MB-231 cells, n = 10, *$p< 0.05$, **$p< 0.01$, ****$p< 0.0001$.

#### Supplementary Fig. S10

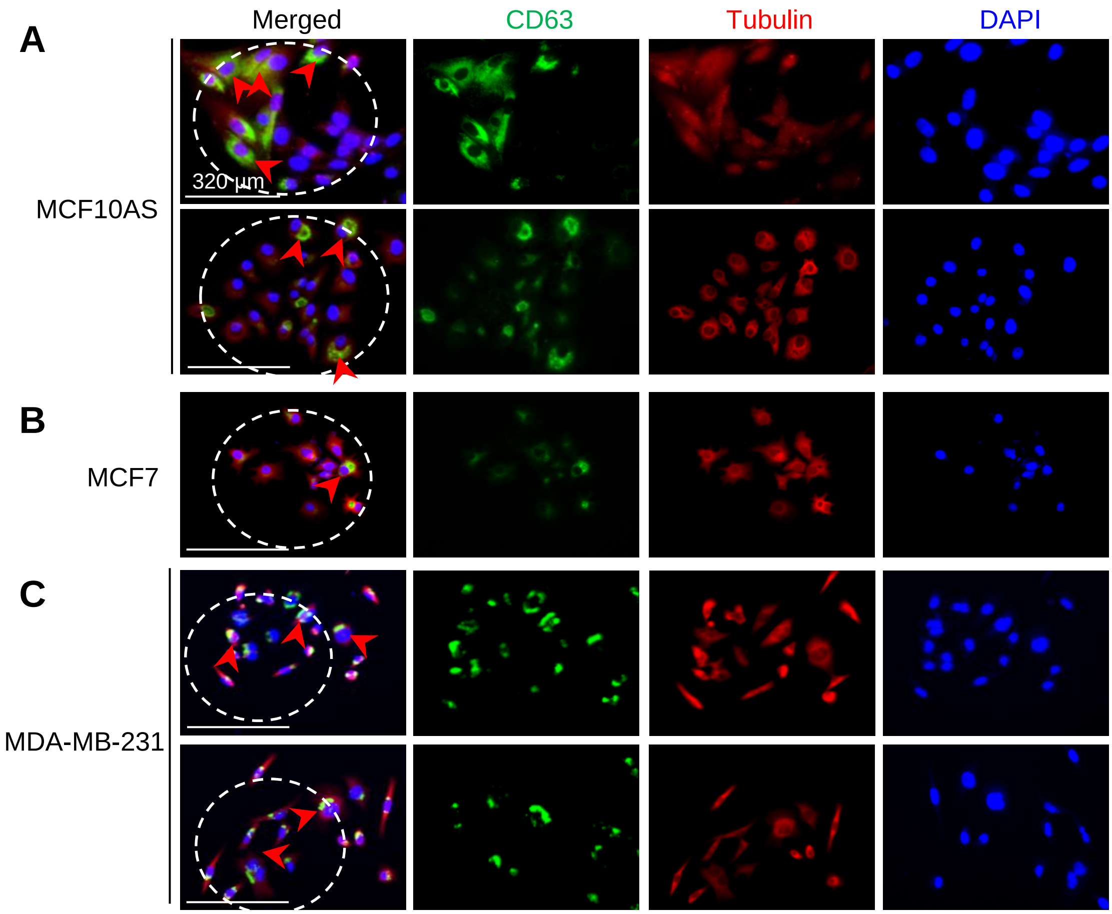


**Supplementary Fig. S10.** PGCs in clonogenic MCF-10AS compared PGCCs in clonogenic BC cells, mouse spontaneous breast tumor and breast cancer tissues. Representative IF images of CD63-expressing PGCs (red arrows) in colonies generated from MCF-10AS (**A**), MCF7 (**B**) and MDA-MB-231 (**C**) cells.
